# Systematic Metaproteomics Mapping Reveals Functional and Ecological Landscapes of Ex Vivo Human Gut Microbiota Responses to Therapeutic Drugs

**DOI:** 10.1101/2025.02.13.637346

**Authors:** Leyuan Li, Caitlin M.A. Simopoulos, Janice Mayne, Zhibin Ning, Xin Zhang, Mona Hamada, James Butcher, Joeselle M. Serrana, Luman Wang, Kai Cheng, Hongye Qin, Krystal Walker, Xu Zhang, Alain Stintzi, Daniel Figeys

## Abstract

Therapeutic compounds exert impacts on gut microbiota; however, how they affect the community functional ecology, especially as reflected at the protein level, remains largely unexplored. In this study, we systematically map metaproteomic responses of *ex vivo* human gut microbiota to 312 compounds, generating 4.6 million microbial protein responses, available as an interactive resource (https://shiny.imetalab.ca/MPR_Viz/). Protein-level analyses identify significant metaproteomic shifts induced by 47 compounds, with neuropharmaceuticals as the sole drug class significantly enriched among these hits. Further analyses on the community level reveal a tri-stability pattern in microbial composition and the emergence of three distinct functional states, based on a functional beta-diversity metric. Notably, neuropharmaceuticals cause particularly strong effects on the microbiomes, lowering the proteome-level functional redundancy and raising the level of antimicrobial resistance proteins, ultimately pushing the microbiome into an alternative functional state. Preliminary validation suggests that enhancing functional redundancy may contribute to maintaining microbiota resilience against neuropharmaceutical-induced antimicrobial resistance. Overall, this work establishes a comprehensive view of how drugs influence gut microbiome function and ecology at the protein level, proposes a landscape-based framework for interpreting community resilience, and highlights the need to consider protein-level and ecological responses in the evaluation of therapeutic interventions.

## INTRODUCTION

The development of human-targeted drugs typically focuses on mechanisms of drug interactions with human protein targets. However, it is often overlooked that therapeutic compounds can have profound impacts on our gut microbiotas, known as off-target effects^1^. Studies have revealed the effects of commonly prescribed drugs on single gut bacterial strains^2^, microbiome composition^3^, function^3,4^ and drug metabolism^5,6^. A more recent study has exemplified these findings by demonstrating that non-antibiotic drugs inhibited *Escherichia coli in vitro* and that the microbial functional pathways they targeted were from those targeted by traditional antibiotics^7^. However, it’s important to note that although *E. coli* serves as a frequently investigated model organism, our gut ecosystem is primarily populated by obligate anaerobes belonging to the Bacillota and Bacteroidota phyla, drug effects on which remain largely unexplored. Additionally, ecological interactions between gut microbial species can lead to more complicated drug responses compared to responses of single species. A microbial community can obtain resistance against drug perturbation through exposure protection^8^ and metabolic cooperation^9^, or conversely, drug-disrupted cross-feeding networks may sensitize the drug response of some microbial species that are otherwise unresponsive to the drug^10,11^. A recent study systematically investigated such community-level behaviors by analyzing species abundance dynamics in a 32-species synthetic community subjected to 30 different drug treatments, and found that 26% of the tested cases exhibited community-level phenomena such as cross-protection and cross-sensitization^12^. Indeed, drugs response involves the complexity of the ecosystem functioning that goes beyond single species survival^13^ or overall compositional responses. In other words, a comprehensive analysis of microbiome drug responses which is observed from a functional omics scope will offer a provide deeper insights of the microbial systems ecology.

The drug modes of action that affect functional microbiome ecosystems vary, but they often involve interactions with key microbial proteins or enzymes. During pharmaceutical drug development, many drugs are developed solely based on their effects through interactions with specific human proteins, without considering the fact that species within the human microbiota may contain structurally similar proteins and may thus also be influenced by these drugs. For example, the serotonin transporter (SERT, encoded by *SLC6A4* in humans), responsible for serotonin uptake in humans, is commonly targeted for pharmaceutical inhibition by neurological drugs such as antidepressants. Interestingly, SERT belongs to the neurotransmitter:sodium symporters (NSS) family, which is conserved from bacteria to humans^14^. Its bacterial homolog, Leucine transporter (LeuT), has been shown to contain a NSS binding site^14^, and has actually been used as a model to predict the antidepressant drug-binding specificity to SERT^15^. Such phenomena suggest that comprehensive observation of drug effects on the metaproteome level – the frontline of microbiota-drug interaction – could offer invaluable insights for in-depth exploration of phenotypic microbiome responses on the functional and ecological levels.

Moreover, since therapeutic drugs span a broad array of chemical structures, they are expected to induce a diverse spectrum of microbial ecosystem responses of the gut microbiota. From the scope of systems ecology, a key question arises: are these responses widely distributed across the functional landscape of the microbiome (a map that connects all possible combinations of microbial functional status)? Or do they cluster around distinct stable functional states, as researchers have suggested regarding community structure^16,17^? Understanding such microbiome functional landscape and mapping drug responses onto it holds significant promise for identifying personalized drug treatments. In microbial systems ecology, functional redundancy (FR) refers to the presence of multiple species that can perform similar functions^18^. It is widely believed that FR is positively related to community resilience^19^. This raises the question of whether FR allows the microbiome to maintain resilience by occupying stable functional states within the functional landscape.

To comprehensively map human gut microbiome’s drug response landscape, robust experimental approaches are fundamentally required to capture a broad range of microbiome functional states within the landscape under diverse drug treatments. *In vitro* microbiome culturing has emerged as a potent tool for uncovering microbial responses to drugs and identify means to manipulate these responses from interference from host factors. We’ve previously developed the RapidAIM platform, which maintains the viability and functional individuality of *ex vivo* human gut microbiota and robustly captures microbiome drug responses using metaproteomics^4^ - a technique for the large-scale identification and quantification of proteins within a microbiome, providing insights into the functional state and activities of the microbial community. We have continued to scale up the RapidAIM workflow, including live microbiota biobanking^20^, cost-effective metaproteomic TMT-labeling^21^, and automation^22^. As a result, we have established the RapidAIM 2.0^22^ platform for high-throughput metaproteomic analysis of *in vitro* microbiota cultures, which was applied in this research to reveal the functional landscape of microbiota drug responses.

In this study, we performed high-throughput assays of 312 commonly prescribed therapeutic compounds against different individual gut microbiotas using the RapidAIM 2.0 platform, resulting in nearly 2,000 microbiota cultures and tandem mass tag (TMT)-labelled metaproteomic screenings. Subsequent in-depth label-free metaproteomics analysis of 110 compounds that passed the initial screening further detailed over 4.6 million bacterial protein-drug responses, provided as a resource through an interactive Shiny app. We analyzed the functional and ecological landscapes of gut microbiota responses to therapeutic drugs across three hierarchical levels: protein-level, revealing metaproteomic shifts in 47 of the tested compounds with neuropharmaceuticals significantly enriched; taxonomic composition-level, exhibiting a compositional tri-stability landscape; and systems ecological-level, where our recently developed PhyloFunc beta-diveristy metric^23^ identified three distinct functional community state types in response to different compounds, demonstrating remarkable systems-level ecological consistency in drug responses across individuals. Notably, specific human-targeted compounds, particularly neuropharmaceuticals, stimulated the expression of microbial antibiotic resistance proteins (ARP), while reducing community-level FR. Building on the functional energy landscape hypothesis, we were able to reduce the neuropharmaceutical-induced ARM increase by introducing a prebiotic into the culture to counteract the FR decrease. These findings underscore the critical need to map the metaproteomic drug response landscape, as it provides a foundational framework for identifying key leverage points to tailor and optimize personalized drug response modulation.

## RESULTS

### The human gut metaproteome is altered by a diverse array of drugs

In this study, we cultured human gut microbiotas from healthy subjects and exposed them to a collection of therapeutic compounds in our RapidAIM assay. The functional responses were measured using high-throughput metaproteomics (Figure 1a). Briefly, the compounds were selected through a multi-step process: we first included 141 drugs with reported impacts on the microbiome based on our in-house database (Supplementary Data S1). Subsequently, we cross-referenced this list with the top 200 most prescribed compounds in the US according to literature^24^, resulting in the addition of 119 compounds. To ensure comprehensive coverage of drug classes and properties, we expanded the list further with 52 additional compounds by addressing three additional questions: 1) according to the ATC classification system, are drug classes being adequately presented or further expansion of drug representatives is needed? 2) does each ATC class contain representative compounds of similar and different chemical structures? 3) which drugs are stereoisomers and can be used to test the stereospecific bioactivity and toxicity? Details of the resulting 312 compounds are listed in Supplementary Data S1. Assessing the compounds using structure-based clustering suggests a good coverage of drug diversity (Figure 1b).

**Figure 1.**
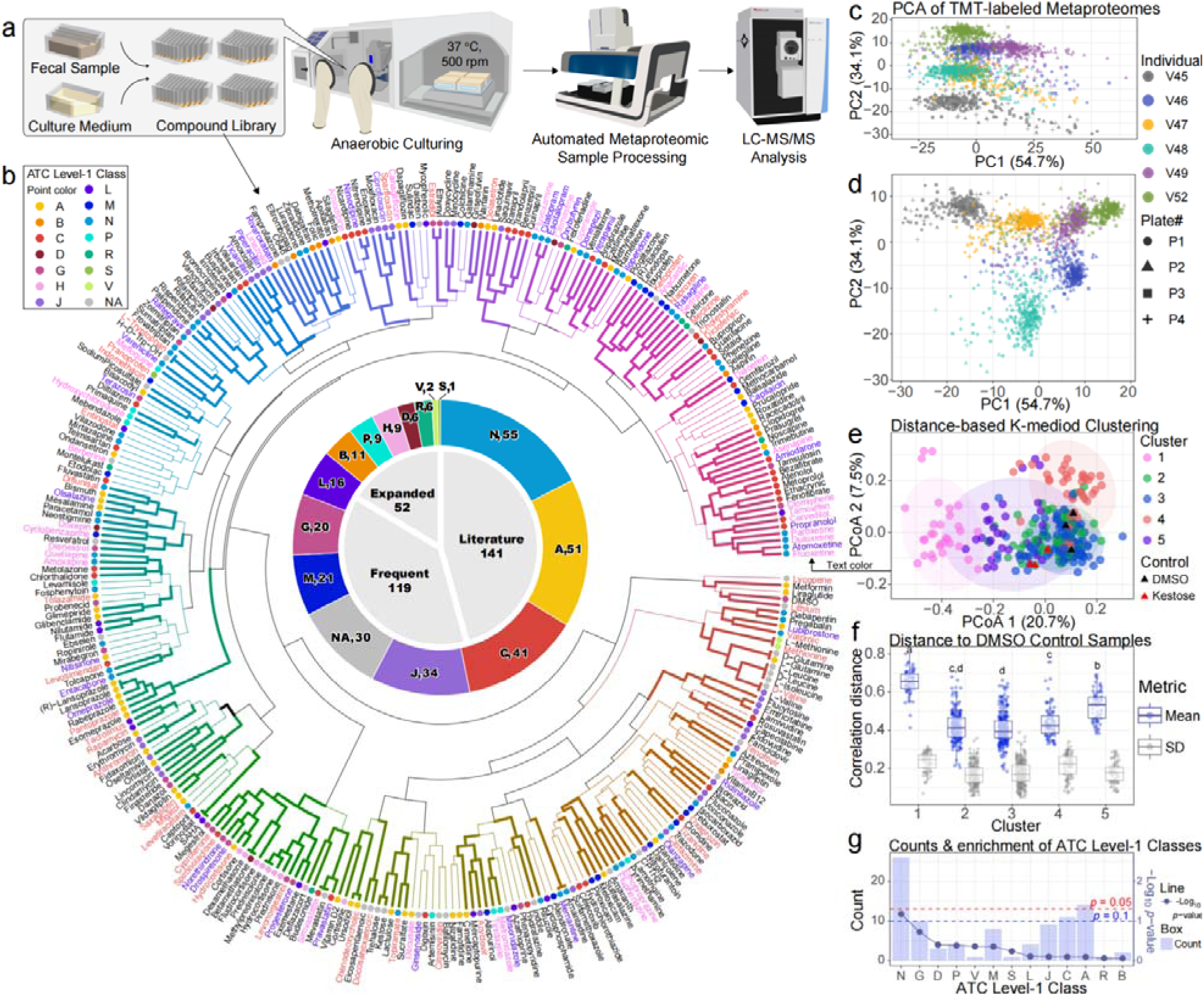
Screening for compounds influencing the gut metaproteome. **(a)** Schematic representation of the experimental workflow. The study follows the RapidAIM 2.0 procedure^22^; fecal slurry samples of each human gut microbiota were cultured anaerobically with the presence of a compound or a control substance (DMSO or kestose). Automated metaproteomics sample processing was performed before samples are analyzed using LC-MS/MS. This figure is adapted from our previously published work^22^ under a Creative Commons Attribution 4.0 International License (CC BY 4.0). (b) Selection, Anatomical Therapeutic Chemical (ATC) classification and structure-based clustering of compounds in the compound library used in this study. Colors of compound labels correspond to three of the clusters in Panel E: magenta – Cluster 1, red – Cluster 4, purple – Cluster 5. Thick branches indicate clusters with high bootstrap support (approximately unbiased (AU) p-value > 95%). (c)-(d) Principal component analysis (PCA) of all samples showing metaproteome individuality as well as responses to compounds. (e) Distance-based K-medoids clustering of all compounds based on the average of six individual microbiome responses. Each point represents a compound or a control. Ellipses represent 95% confidence of each cluster. (f) Distance of compounds to DMSO control samples within each cluster identified in panel (d). The numbers of compound–control pairs were 96, 269, 373, 120, and 99 for clusters 1, 2, 3, 4, and 5, respectively. Mean values and standard deviations were calculated across N = 6 individual microbiome samples. Letters a, b, c, and d indicate levels of significance determined by ANOVA with Tukey’s post hoc test. The center line of box represents the median; the box limits indicate the upper and lower quartiles (Q3 and Q1); the whiskers extend to the most extreme values within 1.5× the interquartile range (IQR) from the quartiles. (g) Enrichment of various ATC level 1 drug classes in Clusters 1, 4, and 5 compounds (corresponding to the clusters identified in panel (d)), among all selected compounds. A one-sided Fisher’s exact test was applied to evaluate whether a given drug class was over-represented in the test set compared to the background, shown are raw *p*-values uncorrected for multiple hypothesis testing.

Experiments were conducted using biobanked live human gut microbiotas from six distinct individuals, with each microbiota treated with different compounds in separate wells of 96-well plates, alongside positive and blank controls (2 mg/mL Kestose and the drug solvent DMSO), resulting in a total of 1,920 microbiota cultures. TMT-11plex^TM^ was used for a first-pass screening of compounds that potentially induced functional responses in the microbiomes. A total of 91,641 peptides (average of 10,025 ± 1,841 peptides per multiplexed TMT set) and 21,859 proteinGroups (average of 5,496 ± 519 proteinGroups per multiplexed TMT set) were identified from the 192 multiplexed TMT sets of 1920 samples. Overall Principal Component Analysis (PCA) based on proteinGroups showed both inter-individual differences in metaproteomics and the alteration of metaproteomes by a subset of compounds (Figure 1c-d). Due to the presence of inter-individual differences, we applied a data analysis pipeline to cluster drug responses by average distances. Briefly, the distance matrix (measured by Pearson’s correlation) between metaproteomic proteome content networks (PCNs)^25^ was first calculated separately for each individual subject; subsequently, the distances between all compounds and controls were averaged across all subjects. Finally, K-medoids analysis revealed five clusters of drug responses (Figure 1e). We showed that three of the compound clusters (clusters 1, 4 and 5) deviated from the controls (Figure 1f). Interestingly, there appeared to be no visible correlation between the drug structure clusters in Figure 1B and the compound response clusters in Figure 1E. From the TMT-based compound response clustering, we show that across Anatomical Therapeutic Chemical (ATC) classification system level 1 classes, the proportion of compounds belonging to each K-medoids clusters are different, we therefore evaluated the enrichment of K-medoids clusters 1, 4 and 5 compounds in each ATC level 1 class. The analysis revealed no significant enrichment for any specific ATC level 1 class compared to the background of all selected compounds. However, it is still notable that Class N (targeting nervous system) showed a *p*-value of less than 0.1 (Figure 1g).

### Comprehensive profiling of metaproteomics drug responses reveals alterations in proteins and functional pathways

A total of 110 compounds from the K-medoids clusters 1, 4 and 5 (Figure 1e and Supplementary Data S1) were selected for further in-depth label-free quantification (LFQ) – based metaproteomics analysis. We used deep metagenomic sequencing of each individual microbiome sample to construct a sample-specific database for the label-free database search following the Metapro-IQ workflow^26^ (Supplementary Figure S1a). A total of 9,326,339 LC-MS/MS spectra were identified as 195,686 unique peptides, corresponding to 36,075 unique proteinGroups. An average of 9,871 ± 3,679 peptides per sample was identified by MS/MS; in addition, 9,994 ± 2,052 proteinGroups were quantified per sample with match-between-runs (MBR) option enabled during protein quantification. The MBR approach allows transfer of peptide identifications across runs based on retention time and mass-to-charge (m/z) alignment, resulting in a higher number of proteinGroups being quantified than peptides directly identified in individual samples. Label-free quantification was then performed using directLFQ^27^ followed by data harmonization using HarmonizR^28^. The processed dataset thus includes over 4.6 million bacterial protein-compound responses. A series of quality-control assessment suggested high reproducibility and robustness of the whole process (Supplementary Figure S1b-d).

Given the extensive spectrum of metaproteomic responses across various functional pathways and taxonomic units, microbial drug responses provide valuable insights into resistance mechanisms, metabolic adaptations, and ecological interactions under therapeutic interventions. To facilitate data exploration and hypothesis generation in future drug-microbiome research, we have developed an interactive Shiny app as a resource for exploring functional and taxonomic responses to these 110 compounds, available at https://shiny.imetalab.ca/MPR_Viz/. This app allows readers to select specific ATC drug classes and explore the functional and taxonomic responses of individual compounds within these classes. In more details, the features that can be visualized involve functional annotations such as NOG categories and NOG accessions (annotated by eggNOG mapper), KEGG KO annotations (from the KEGG database), as well as protein abundances across different taxonomic units. By providing this level of detail, the app facilitates a comprehensive exploration of the drug-microbiome interactions at both the functional and taxonomic levels. Readers can visualize data and download the figures, making it a highly informative tool for further exploration and hypothesis generation about microbiome responses to drugs. Here, we show an example of the Shiny app interface with an analysis result of showing several significant increases in the NOG annotation of proteins associated with efflux transmembrane transporter activity following independent treatments of several different drug from the ATC Level 1 class N (Supplementary Figure S2). This suggests that exposure to these drugs may have stimulated an increase in drug resistance activity, likely through the upregulation of efflux transporters.

We next examined the protein-level responses by performing PCA on the label-free quantified proteinGroup intensities using the nonlinear Iterative Partial Least Squares (NIPALS) algorithm^29^. As shown in Figure 2A, the PCA plot reveals a densely clustered distribution due to the large number of samples. Nonetheless, a global shift from the control group is evident, indicating widespread compound-induced alterations in the metaproteomic profiles of the selected compounds. We further examined the PCA within each distinct ATC class and at various ATC levels (Supplementary Figures S3-S4). While only a subset of compounds in each of ATC level-1 classes A (alimentary tract and metabolism), B (blood and blood forming organs), D (dermatologicals), L (antineoplastic and immunomodulating agents), and M (musculo-skeletal system) showed separation from the controls, all compounds in ATC level-1 class P (antiparasitic products, insecticides and repellents) separated from the controls on PC 1 which explains 62.7% of the variance. Classes C (cardiovascular system), G (genito urinary system and sex hormones), J (antiinfectives for systemic use) and N (nervous system) contained a larger number of compounds. We performed further explorations of PCAs on the second ATC level and presented classes that contained discriminative compounds (Supplementary Figures S3). On ATC level-2, we show that classes G03 (sex hormones and modulators of the genital system), J01 (antibacterials for systemic use), N04 (anti-parkinson drugs), N05 (psycholeptics) and N06 (psychoanaleptics) classes contained greater than three compounds discriminated from the controls. Among these, the ATC level-4 code N06AB (selective serotonin reuptake inhibitors) comprised six compounds with noticeable effects (Supplementary Figures S4). By further examining PCA by individual compounds, we observed a total of 47 compounds that are discriminative from the control in all individual microbiomes, denoted as Metaproteomics-response positive (MPR+) drugs (Supplementary Data S2 and Figure 2b). Among which, 39% are Class N compounds, and 15% are Class C compounds. Enrichment analysis of these 47 compounds against the background of all 312 drugs showed a significant enrichment of Class N drugs (Figure 2c), suggesting that drugs targeting the nervous system have the most frequent off-target effects on gut microbes.

**Figure 2.**
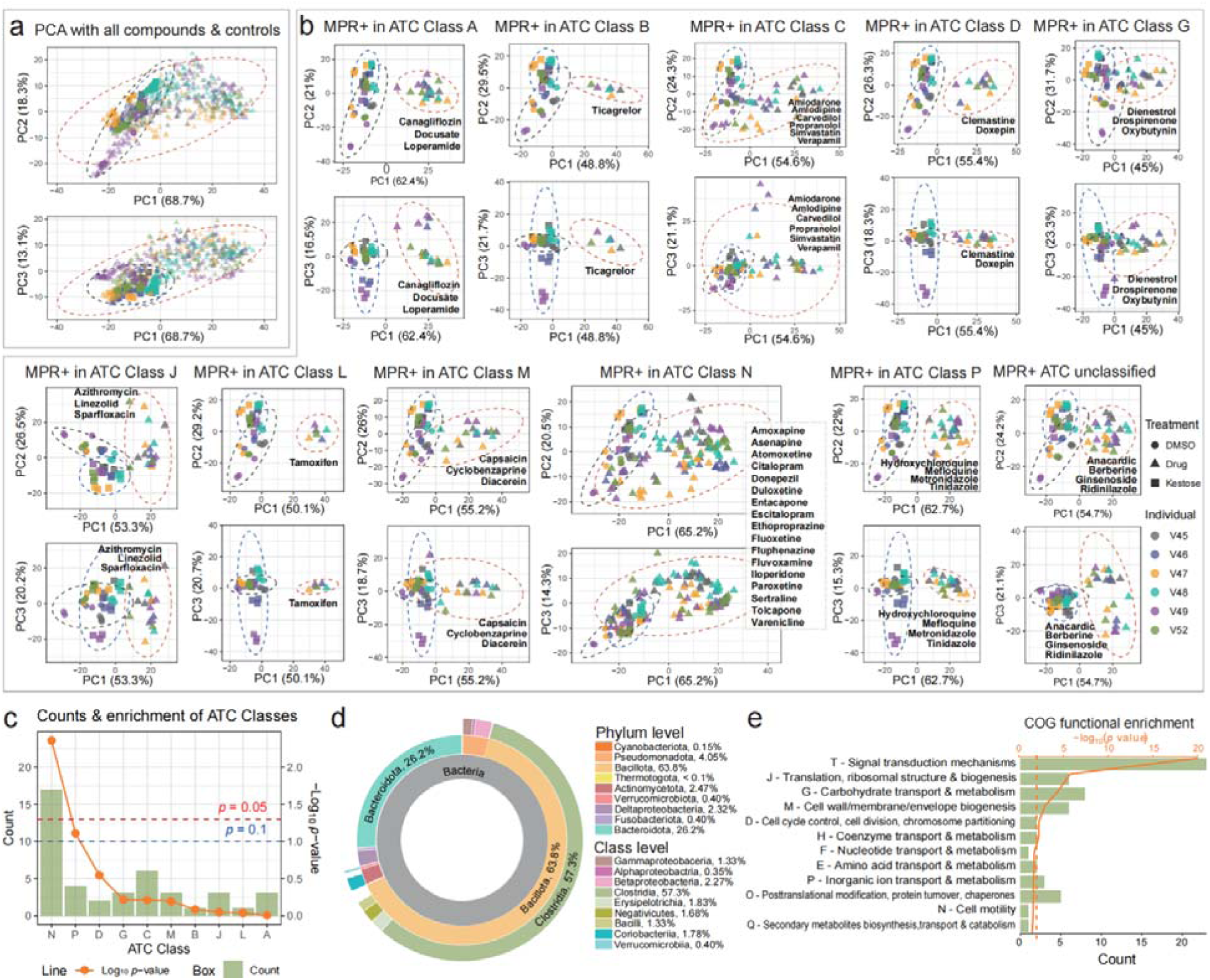
Discovery of MPR+ compounds and microbial homologs of their human targets. (a) Overall PCA of the label-free dataset. (b) Principal component analysis (PCA) of metaproteomics response-positive (MPR+) compounds and control samples, grouped by ATC level-1 classification. For (A)-(B), the first three components of Principal Component Analysis (PCA) are shown. All six individual gut microbiome are shown as labeled by different colors. Gray dashed ellipse denotes 95% confidence interval of each group (red: drugs, black: DMSO and blue: kestose). (c) Enrichment analysis of ATC level 1 class in MPR+ compounds against all 312 selected compounds. A one-sided Fisher’s exact test was applied to evaluate whether a given drug class was over-represented in the test set compared to the background, shown are raw p-values uncorrected for multiple hypothesis testing. (d) Discovery of 2,528 microbial proteins homologous to human drug targets in the individual gut microbiomes’ metaproteomics database using the HMMER algorithm. The homologous proteins were annotated by eggNOG mapper, and their taxon-specific annotations were summarized and visualized in a Circos plot. (e) COG functional enrichment of the identified homologs was then performed using iMetaShiny, in which a one-sided hypergeometric test was used.

We were curious about which human proteins are targeted by these MPR+ compounds and whether these targets could have bacterial homologs. Therefore, by searching MPR+ compounds against the molecular drug target database by Santos et al.^30^, we extracted a list of 50 human protein targets (Supplementary Data S2). KEGG enrichment analysis shows that these proteins are most significantly enriched in neuroactive ligand-receptor interaction and calcium signaling pathways (Supplementary Figure S5a-b), both are targets of Class N drugs. Using the HMMER algorithm^31^ to map the drug target Pfams through our metaproteomic sample-specific database, we discovered 2,528 microbial proteins that are homologous to these human targets. EggNOG annotation was able to map 2,115 of these proteins to reference taxa and functions which were mainly originated from the Clostridia class, with signal transduction mechanisms being the most significantly enriched functional category (Figure 2d-e, and Supplementary Data S3). Notably, 270 bacterial proteins homologous to human neurotransmitter sodium symporters, the targets of selective serotonin reuptake inhibitors (SSRIs) and serotonin-norepinephrine reuptake inhibitors (SNRIs), were identified within the sodium:neurotransmitter symporter or sodium:solute symporter family.

### Taxonomic responses to drugs exhibit a compositional tri-stability landscape

Taxonomic responses, as it takes account into cumulative abundances of taxon-specific peptides, provides an alternative view of microbiomes community responses that is different from protein-level PCA which takes into consideration proteins as features. The compounds selected for LFQ analysis span a wide spectrum of drug structure and ATC classes. Our initial intuition was that the microbiome responses to different compounds can be highly diverse, resulting in a diverse combination of microbiome composition. The equal-volume processing protocol (see Methods > RapidAIM 2.0 assay) ensures direct comparability of protein-based taxonomic abundances between samples. To examine the microbiome responses, we selected the microbial genera that showed at least one significant drug response across all six microbiomes, and visualized their abundance changes in a genus-level heatmap (Figure 3A). Despite observing highly diverse patterns in protein-level analysis, we identified an intriguing phenomenon in the taxonomic (genus-level) response heatmap: there’s a coherent transition between three clusters of responses (Figure 3a). Among which, one cluster (Cluster 2) only included five genus-level alterations, while the two other clusters (Clusters 1 and 3) exhibited more extensive changes in 44 genera in response to 63 compounds. By color-coding different genera in each cluster with blue, orange, and green tones, the overall transition becomes apparent. This can be more easily observed by visualizing the drug response spectrum of each individual microbiome through stacked column bar plots (Figure 3b).

**Figure 3.**
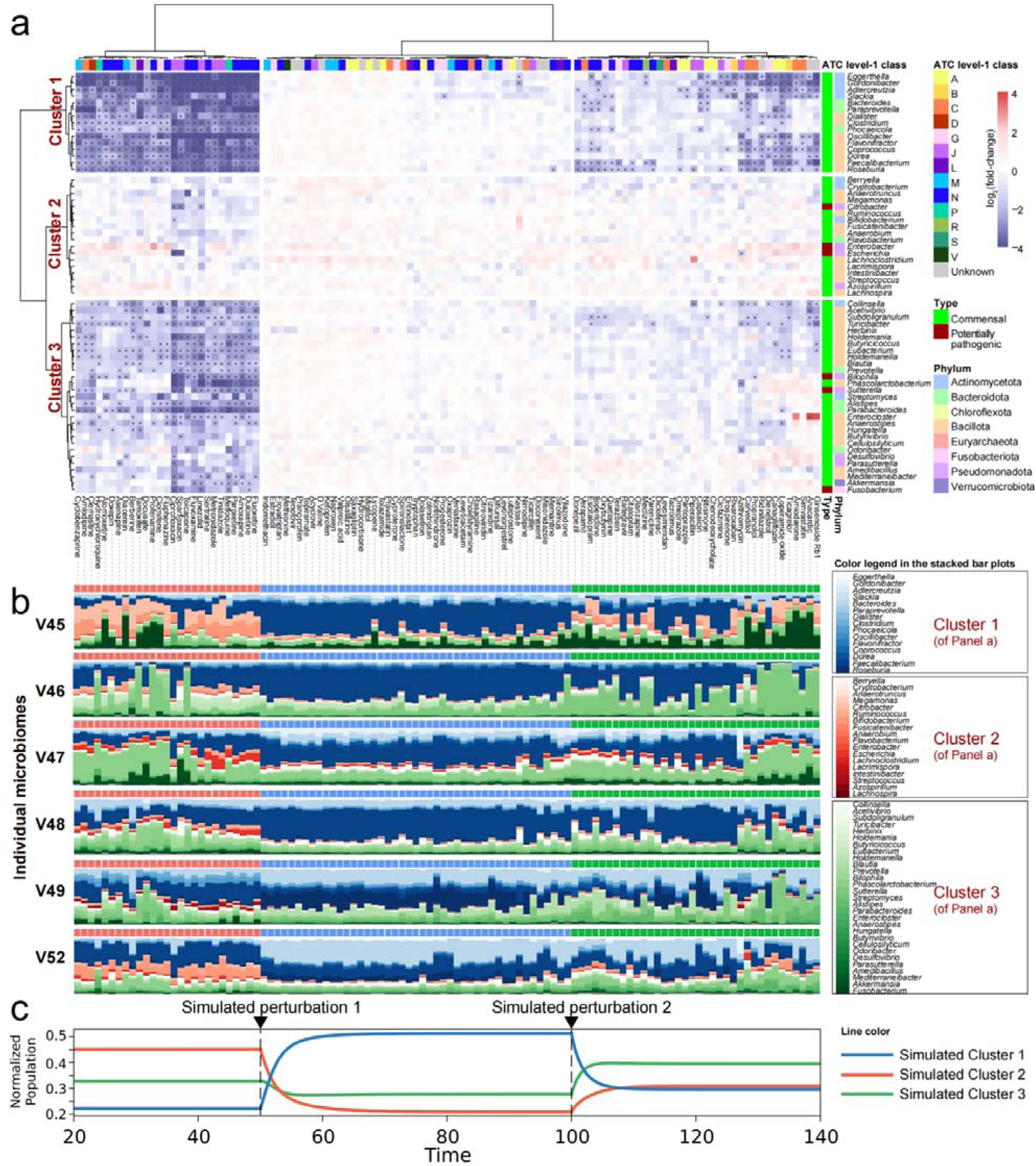
Taxonomic responses reveal multi-stability of individual gut microbiomes driven by drug stimulation. (a) Heatmap and hierarchical clustering showing log_2_ fold-change of overall protein biomass specific to different genera. Asterisks indicate significance levels from a two-sided Wilcoxon signed-rank test, with *p* < 0.05 and fold-change >1.5. For each genus-compound pair, the test was performed to evaluate whether the distribution of values significantly deviated from the reference value of 1, under the null hypothesis that the median of the differences (Value − 1) is zero. A continuity correction was applied to the test statistic to improve the normal approximation. (b) Relative abundance of genus-specific proteins presented in stacked column bar plots. The sample order agrees with panel A, and different genera are also colored according to the hierarchical clustering in panel A. (c) gLV competition model considering between-cluster interactions simulates the switches between three different stable states under drug perturbation. The time axis in panel C is conceptual and does not correspond to the actual order of drug treatments depicted in panels A and B.

This provides evidence for the theory of multiple stability points in the microbiome^16,32^, our work marks the first observation of a tri-stability landscape of individual microbiome compositions within the drug response space. To explain the phenomenon of steady states in the functional response of the microbiome to drugs, we used the generalized Lotka-Volterra (gLV) to describe the response dynamics (see Methods). In this model, we classified the genera observed in the experiment into three distinct clusters based on co-abundance patterns, assuming that interactions within each cluster are negligible. By adjusting the model parameters to simulate the effect of drug stimulation, we can reproduce the switches from one stable state to another (Figure 3c), mirroring the actual microbiome responses observed in Figure 3b.

### Community state analysis reveals universal drug-specific microbial ecological responses

Examining protein and taxonomic compositions separately fails to capture the full ecological landscape from a systems perspective. Therefore, it is crucial to adopt a more ecologically relevant assessment of microbiome responses. We have previously developed a tool named PhyloFunc, which integrates phylogenetic and functional information to assess microbial community dissimilarities (i.e. beta diversity) on the ecosystem level^23^. PhyloFunc analysis was performed on each individual’s microbiome including all LFQ samples of drug treatment and control conditions. The PhyloFunc distance results from different individuals were combined and visualized using Principal Coordinates Analysis (PCoA). K-means clustering was then applied, identifying three distinct community-state types (CSTs) (Figure 4a), with the optimal number of clusters determined as three based on the Gap statistic (Figure 4b). Based on these three clusters, we used a Sankey diagram to visualize the distribution of drug names and their sources across different CSTs (Figure 4c). Notably, all negative control (DMSO) individual microbiomes clustered into CST-3, while 41 of the compounds caused at least 80% of the tested individual microbiomes to shift into CST-1 and CST-2. These compounds include Amiodarone, Amlodipine, Amoxapine, Anacardic, Asenapine, Atomoxetine, Berberine, Canagliflozin, Carvedilol, Clemastine, Clomiphene, Cyclobenzaprine, Diacerein, Dienestrol, Docusate, Doxepin, Drospirenone, Duloxetine, Entacapone, Escitalopram, Ethoproprazine, Fluoxetine, Fluphenazine, Fluvoxamine, Ginsenoside, Hydroxychloroquine, Iloperidone, Linezolid, Loperamide, Mefloquine, Metronidazole, Oxybutynin, Paroxetine, Ridinilazole, Sertraline, Simvastatin, Tamoxifen, Ticagrelor, Tinidazole, Tolcapone, and Verapamil.

**Figure 4.**
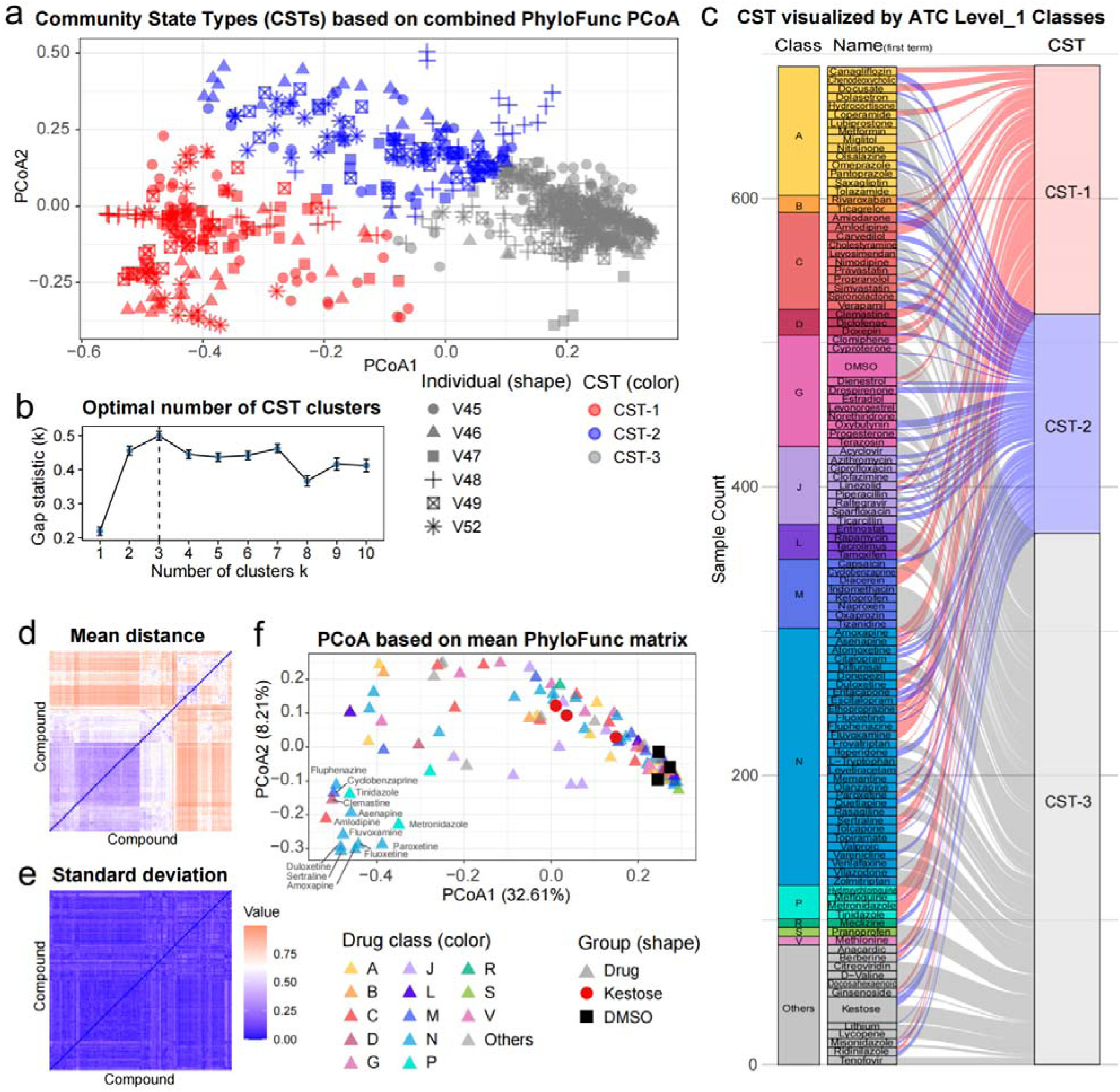
PhyloFunc analysis reveals impact of drugs on microbial community states. (a) Principal Coordinates Analysis (PCoA) of PhyloFunc distances, clustering microbiome responses into three distinct Community-State Types (CSTs). (b) Determination of the optimal number of CST clusters using the Gap statistic ( , represented by centerlines). Error bars represent the standard error ( ) values across Monte-Carlo reference simulations at each . (c) Sankey diagram illustrating the distribution of drugs across CSTs, showing the association of compound names and their ATC level-1 classes with different CSTs. (d) Heatmap of the Mean PhyloFunc distance matrix, representing the averaged PhyloFunc distance for each compound-compound pair across individuals (N = 6). (e) Heatmap of the Standard Deviation (SD) of PhyloFunc distances (N = 6). (f) PCoA based on averaged PhyloFunc distance matrix across all individuals.

To further illustrate that PhyloFunc’s inter-individual variability is not affected by individual-specific differences in protein composition, we calculated the Mean and standard deviation (SD) of PhyloFunc distances across individuals (N = 6) (Figure 4d-4e). We can clearly observe the overall differences in heatmap intensity between the Mean and SD matrices when using the same color gradient scale. To further quantify these differences, we applied a two-sided Wilcoxon signed-rank test between the Mean and SD, which demonstrated a highly significant difference (p-value < 2.2e-16, N = 12,321 compound-compound interactions). Such systems-level ecological consistency in drug responses across individuals enabled us to summarize the PhyloFunc results into an averaged distance matrix (Figure 4f), where we identified a distinct cluster of 13 drugs, 8 of which belonged to the N-class category

### A subset of neuropharmaceuticals exerted similar functional effects on the ex vivo human gut microbiomes

As an example of specific interest in taxonomic and functional responses, we observed that a subset of neurological drugs exerted nearly identical effects on the human gut microbiomes as assessed by protein-level PCA, including seven drugs, namely amoxapine (ATC N06AA), duloxetine (ATC N06AX), fluoxetine (ATC N06AB), fluphenazine (ATC N05AB), fluvoxamine (ATC N06AB), paroxetine (ATC N06AB), and sertraline (ATC N06AB) (Figure 5a). We show that within each individual microbiome, NIPALS distances between pairs of these compounds are not significantly different compared to that between technical replicates of the DMSO and kestose control groups. In contrast, this group of compounds showed significantly higher distances from both the controls (Figures 5b and 5c). Interestingly, they did not belong to a same ATC level-3/4 class, while most of these drugs are antidepressants that have been reported to have a common target of human SERT proteins, fluphenazine belongs to antipsychotics that does not target SERT. We therefore considered these drugs as a group of potentially common mode of action on the gut microbiome and visualized differentially expressed proteins of this drug group using volcano plot (Figure 5d). In comparison to the DMSO control group, we observed 1,447 significantly decreased proteins and 357 significantly increased proteins. By annotating the proteins to taxon (genus level) and function (COG category), we observed that the functional responses showed strong taxonomic specificity, characterised by overall decreased growth of commensals and enriched proteins of opportunistic pathogens in Pseudomonadota (Figure 5e-f). Taxa having decreased functional protein classes are consistent with previous reports indicating that these taxa are also suppressed^2,33^.

**Figure 5.**
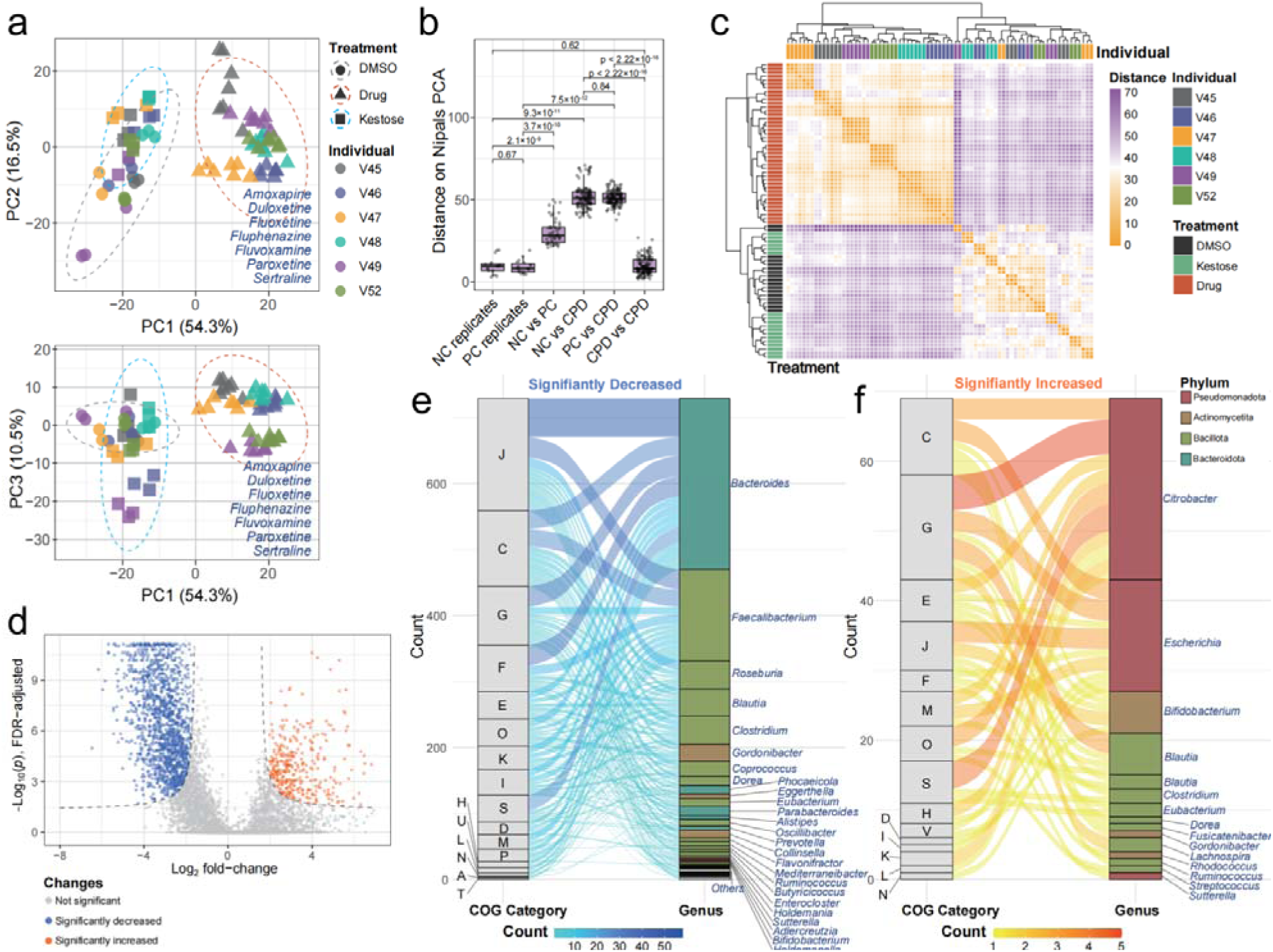
Seven neurological drugs showing near identical metaproteomic effects. (a) NIPALS PCA showing discriminations between the seven selected neurological drugs and the controls. (b) Between-sample distances were calculated based on NIPALS scores and the resulted distances were compared between groups. NC, negative control (DMSO); PC, positive control (kestose); CPD, seven selected compounds. Two-sided Wilcoxon test was applied, **** indicates p < 0.0001. From left to right, the numbers of between-sample comparisons are N = 16, 18, 51, 119, 126, and 126, respectively. The center line of box represents the median; the box limits indicate the upper and lower quartiles (Q3 and Q1); the whiskers extend to the most extreme values within 1.5× the interquartile range (IQR) from the quartiles. (c) The NIPALS score-based distance matrix was further visualized using heatmap and hierarchical clustering. (d) Volcano plot visualizing log2 fold-change of proteins versus FDR-adjusted *p* values (-log_10_) using two-sided Wilcoxon rank sum test. Significantly differential features were determined by log_2_ fold-change threshold of 1.5 and FDR-adjusted *p* value of 0.05. Threshold curves^34^ were defined using a curvature of 1 as previously described. (e, f) Sankey plots showing taxon-specific functional responses of the significantly decreased (e) and increased (f) proteins.

### Antimicrobial resistance negatively correlates to functional redundancy

We have previously performed a proof-of-concept (POC) study of the RapidAIM assay using 43 compounds, and found that drugs not only cause changes in functional and taxonomic composition, but also induce changes each individual microbiomes’ levels of drug resistance proteins^4^. We further wondered whether these changes in predicted drug resistance and alteration of taxonomic composition had any relationship with a community’s functional ecology properties (Supplementary Figure S6a). Recently, we have developed an approach to quantify a microbial community’s proteome-level functional redundancy (*FR_p_*) and its normalized version relative to the community’s taxonomic diversity, *nFRp*^25^ (Supplementary Figure S6b). *FR_p_* describes the ability of multiple taxonomically distinct organisms to contribute in similar ways to an ecosystem, providing informative insights into microbial community ecology on the functional level. Here, we first reanalyzed the RapidAIM POC dataset^4^ and compared *nFR_p_* levels with community composition and levels of antimicrobial resistance proteins (ARP) (Supplementary Figure S6c). We observed instances where increased prevalence of ARP occurred concurrently with a decrease in *nFR_p_* under rifaximin (Supplementary Figure S6D), berberine, flucytosine, and metronidazole treatments. This case study roused our interest, and we were curious whether such phenomenon would be reproduced in this expanded collection of drug-microbiome cultures.

Hence, we analyzed the LFQ dataset using the full CARD antimicrobial resistance database^35^ to redo database search and sum ARP abundance to estimate community-level drug resistance. We observed a strong negative correlation between fold-change of *nFR_p_* and ARP ratio (Pearson’s product-moment correlation coefficient *r* of – 0.695 ± 0.154, Figure 6a). Interestingly, *nFR_p_* response and ARP ratio were both independent of taxonomic diversity (TD) responses (Figures 6b and 6c), suggesting that antimicrobial resistance corresponds to the functional ecology status of a community, irrespective of taxonomic diversity. We also observed a mixture distribution on the scatter plot, which becomes particularly evident when performing K-medoids clustering with each individual microbiome dataset (Figure 6d). This suggests the presence of alternative functional states of the microbiome, wherein the microbiome can transition from its original functional ecological state of high to an alternative state of low when influenced by specific groups of compounds. We demonstrated this concept by illustrating the energy landscape of individual V45’s microbiome using the Kernel Density Estimation (KDE) method (Figure 6e). The upset plot demonstrates a high level of common presence of these compounds across each individual microbiome’s high clusters (Figure 6f). We calculated the frequency of compounds in the high clusters by summing their occurrences across all individual datasets, N05A (antipsychotics) and N06A (antidepressants) drug classes of ATC level 3 showed a high frequency of appearance in the high clusters compared to other N class drugs targeting the neural system (Figures 6g-h).

**Figure 6.**
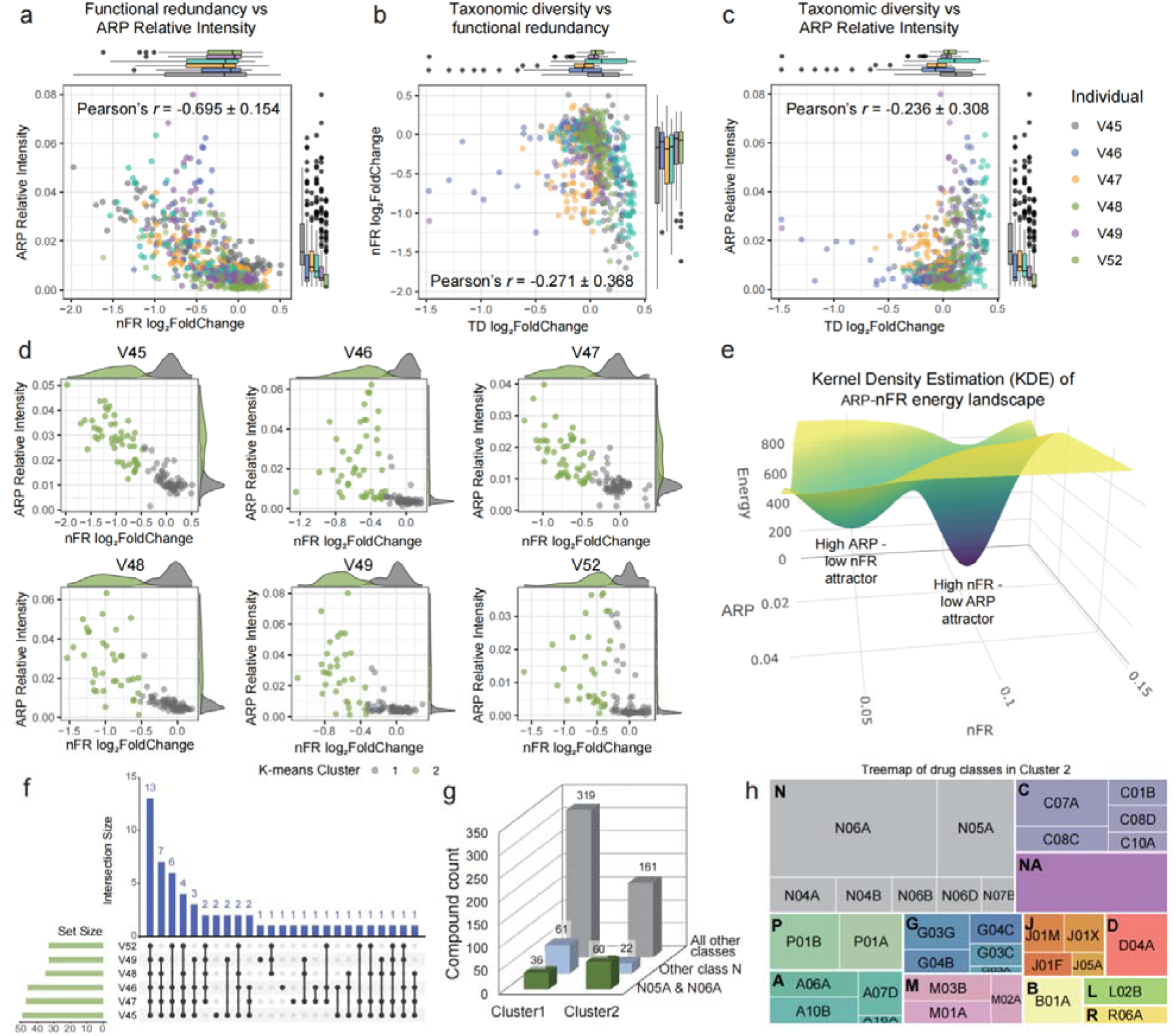
ARP abundance negatively correlates with functional redundancy. (a) Scatter plot and Pearson’s correlation between functional redundancy log2 fold-change and ARP ratio. (b) Scatter plot and Pearson’s correlation between taxonomic diversity log2 fold-change and functional redundancy log2 fold-change. (c) Scatter plot and Pearson’s correlation between taxonomic diversity log2 fold-change and ARP ratio. In panels (a)-(c), box plots show value distributions based on individual microbiome corresponding to each axis. Each box includes N=115∼117 compound-treated or control metaproteomes for each subject’s sample. The thicker black line in the box plot denotes median, the lower and upper hinges correspond to the first and third quartiles, the box ranges from the 1.5× (interquartile range) below the lower hinge to 1.5×IQR above the upper hinge, and whiskers represent the maximum and minimum values, excluding outliers. (d) K-medoids clustering of ARP-FR response by individual. (e) Energy landscape of ARP-nFR response by KDE method. (f) Set comparison of Cluster 2 compounds across individuals, with bar heights representing the number of compounds in each intersection. (g) Counts (represented by bar heights) of Cluster 1 & 2 compound frequencies in different classes. (h) Tree map showing frequency of occurrence in Cluster 2. Bold capital letters indicate ATC level 1 classes, and capital letters with numbers indicate ATC level 3 classes.

### Enhanced functional redundancy mitigated drug-induced ARP

Considering the ARP-*nFRp* drug response energy landscape shown in Figure 6e, and based on the dynamic mechanisms of energy landscapes, we wondered if functional redundancy could be leveraged to prevent drug-induced shifts of the microbiome’s functional state from the high *nFR_p_* - low ARP attractor to the high ARP-low *nFR_p_* attractor. Specifically, our hypothesis was that while a microbiome was stimulated by an ARP-inducing drug (for example, cluster 2 drugs), adding another compound that can increase a microbiome’s functional redundancy may help combat the functional redundancy decline and suppress ARP. To test this hypothesis, we designed a 2x2 factorial design as shown in Figure 7a.

**Figure 7.**
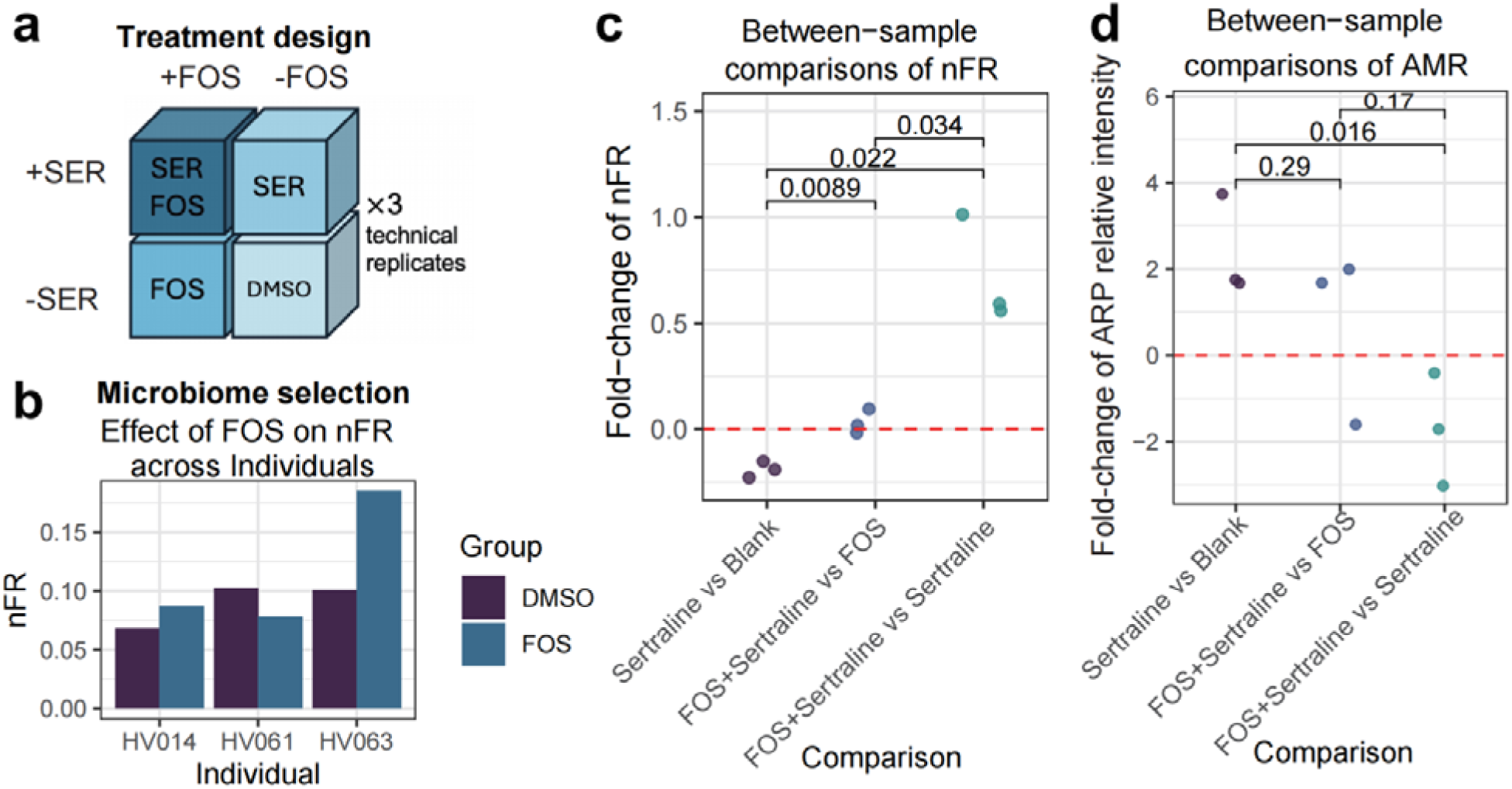
Functional redundancy acts as a potential target to combat drug-induced ARP. (a) Experimental design, sertraline was abbreviated as SER on the figure. (b) Pretest of individual gut microbiome responses to FOS show individualized nFR alterations. One single value is represented by each bar. (c) Between-sample comparisons of nFR and (d) ARP, two-sided t-test was performed.

We selected sertraline as our model for an ARP-inducing drug, as it showed the strongest effect in reducing *nFR_p_* and increasing ARP levels across all six individual microbiomes (Supplementary Figure S7a). Sertraline was also shown to be pathogen-favoring in previous research^36^. Fructooligosaccharide (FOS) was selected as a *nFR_p_* enhancing compound because it increased *nFR_p_* in a subset of previously tested individual microbiomes^4^ (an example is shown in Supplementary Figure S6c). Since the response to FOS can vary between individuals, we performed a pretest using three additional individual microbiomes and selected one microbiome with the highest level of increase in the presence of 10 mg/mL FOS (Figure 7b) and performed the 2x2 factorial design experiment. The effects of the different combinations on and ARP are measured using metaproteomics. Results show that sertraline alone decreased nFR and increased ARP (first group of comparisons in Figure 7c and d). When sertraline was combined with FOS, it did not decrease nFR nor increase ARP, as compared to the microbiome treated by FOS alone (second group of comparisons in Figure 7c and d). Furthermore, compared to the microbiome treated only with sertraline, those treated with both sertraline and FOS showed significantly increased and decreased ARP (third group of comparison in Figure 7c and d). In addition, sertraline increased the abundance of Enterobacterales, but this increase did not occur when sertraline was present alongside FOS (Supplementary Figure S7b). These findings together suggest the possibility that manipulating the microbiome to increase may help counteract the increase in ARP and contribute to preserving microbiome resilience against drug-induced negative disruptions. However, due to inter-individual variability, future studies in larger populations are needed to assess the generalizability of this strategy.

## DISCUSSION

In this study, we systematically profiled how 312 commonly prescribed and structurally diverse drugs and compounds perturb human gut microbiomes ex vivo using the RapidAIM 2.0 platform. Based on an initial TMT-based screening, 110 drugs that induced distinct microbiome perturbations were selected for in-depth label-free quantitative (LFQ) metaproteomic analysis across the microbiomes. The study was adequately powered for exploratory discovery, enabling the detection of key ecological patterns in drug–microbiome interactions. A central finding of this work is that gut microbiome responses to drugs exhibit a reproducible pattern at the ecological systems level, even amid substantial inter-individual variability in baseline microbiome profiles, and that these responses can potentially be modulated by leveraging functional redundancy as an ecological leverage point.

At the functional and taxonomic levels, we found that drugs targeting the nervous system (ATC level-1 class N), cardiovascular system (C), genitourinary system (G), and anti-infectives (J) induced the most pronounced metaproteomic shifts. Among these, nervous system drugs— including SSRIs and SNRIs—consistently elicited strong perturbations in microbial protein expression and community composition. This was consistent with previous findings that SSRI treatment alters gut microbiota composition and function in humans and animal models^37–39^. Our analysis suggests that these effects may involve unintended interactions between drugs and bacterial homologs of human protein targets. Specifically, we observed over 200 protein-coding genes from the sodium:neurotransmitter symporter or sodium:solute symporter family in the tested gut microbiomes. These bacterial transporters are homologous to human neurotransmitter sodium symporters (NSS), which are the targets of commonly used antidepressants such as SSRIs and SNRIs, including the human serotonin transporter (SERT)^30,33^ responsible for serotonin reuptake. However, although these proteins were detected during the first search in our two-step metaproteomic workflow^26^, they were not consistently quantified in the second search stage, likely reflecting their relatively low abundance and representing a limitation of current metaproteomic approaches, which constrained our ability to quantify the responses of these proteins to the neurological drugs. Another limitation of this study is that we did not directly assess the interactions between drugs and bacterial homologs of human protein targets in the human gut microbiomes. Establishing such molecular-level evidence will be critical for future investigations to fully elucidate the mechanisms underlying drug–microbiome interactions.

We next explored the ecological responses at the taxonomic composition level. Counterintuitively, the microbial community composition did not exhibit a broad continuum of distributions but instead formed three distinct states, suggesting the presence of intrinsic multi-stability in gut microbial communities. To further investigate the theoretical principles underlying this tri-stability, we applied the generalized Lotka-Volterra (gLV) model to describe the response dynamics and understand the stability of microbial community states. However, we acknowledge that our current model is theoretical and may oversimplify the actual microbiome dynamics. Moreover, while our findings highlight interesting patterns in taxonomic responses, taxonomic analyses alone are insufficient to fully reveal the ecological response landscape of the microbiome, as they do not adequately capture how microbial functions reorganize in response to drug perturbations—especially given the extensive functional redundancy within microbiomes.

By contrast, metaproteomic data provide a valuable foundation for systems-level ecological analysis, as they directly reflect the functional state of the community under perturbation. To leverage this potential, we applied our previously developed *PhyloFunc* framework, which integrates phylogenetic relationships and functional annotations to quantify ecological distances between microbiome states. Unlike protein-level PCA approaches that rely on data harmonization methods to minimize inter-individual protein variability, PhyloFunc captures ecosystem diversity directly from the original protein group data without requiring data harmonization across individuals. It reduces the impact of compositional differences among phylogenetically related species, thereby offering a more ecologically meaningful perspective on functional community responses. Our finding suggests that while metagenomics and metaproteomics have traditionally revealed high inter-individual microbiome diversities in strains, genes, and proteins, a notable level of consistency emerges at the functional ecosystem level. This suggests that drugs may induce conserved shifts in the ecological function of gut microbiomes across individuals—an insight that cannot be fully captured by taxonomic or protein-level analyses alone. Our findings emphasize that integrating functional and taxonomic dimensions through an ecological framework is essential for understanding drug–microbiome interactions.

Notably, in all different levels of response landscapes, we observed that several drugs targeting the nervous system -amoxapine, duloxetine, fluoxetine, fluphenazine, fluvoxamine, paroxetine, and sertraline —consistently induced strong perturbations in gut microbial proteins and community ecology. These drug effects were associated with decreased commensal growth and enrichment of pathogen-associated proteins, raising concerns about their long-term impact on gut ecosystem health. The complexity of the microbiome makes it unlikely that off-target drug effects can be mitigated by targeting individual species. Instead, our findings underscore the need to consider the functional ecological landscape of the microbiome as a framework for understanding and managing drug-microbiome interactions. Emerging from our analyses was the observation of bimodal functional states within the microbiome drug response landscape: a high FR ^25^, low ARP state, and a low *FR_p_*, high ARP state. These patterns resemble ecosystem regime shifts described in resilience and redundancy theory^40^, wherein loss of functional redundancy may lower the resilience of the microbial community, increasing susceptibility to dominance by resistant strains. Notably, such non-linear dynamics are not readily captured by taxonomic or single-protein analyses alone but require a systems-level ecological perspective. We show that by using prebiotic FOS, the ecosystem can potentially buffer against the functional redundancy loss in specific individual microbiomes and prevent the system from shifting to an undesirable state dominated by resistant strains. However, further validation in larger and more diverse cohorts will be essential to translate these insights into personalized strategies for maintaining gut ecosystem health during pharmacological treatments.

Several limitations of this study should be acknowledged. First, as an ex vivo microbiota culture model, the RapidAIM 2.0 platform cannot recapitulate host–microbiome interactions; rather, it enables the investigation of microbiome-intrinsic responses to drugs, independent of host factors. Second, as discussed earlier, this study did not directly investigate the molecular mechanisms by which drugs interact with microbial proteins, though our findings highlight their potential importance in shaping drug–microbiome dynamics. Looking forward, integrating systems ecology frameworks with mechanistic and longitudinal approaches will provide deeper insights into drug– microbiome interactions and help inform strategies to preserve microbiome resilience and promote recovery from drug-induced perturbations.

## METHODS

### Sample collection and biobanking

The protocol for human stool sample collection (# 20160585–01 H) was approved by the Ottawa Health Science Network Research Ethics Board at the Ottawa Hospital, Ottawa, Canada. The inclusion criteria of participants were 18-65 years of age and healthy. Exclusion criteria included diagnosis of irritable bowel syndrome, inflammatory bowel disease, or diabetes; antibiotic use or gastroenteritis episode three months preceding collection; use of pro-/pre-biotics, laxatives, or anti-diarrheal drugs in the last month preceding collection; or pregnancy. In this study, we collected samples from nine healthy subjects (23-48 years of age, four females and five males). Sex/gender of participants was self-reported. Written informed consent was obtained from all participants for the collection and use of samples and data analysis for the present work. Sex/gender was not considered as a study variable in the experimental design, as the study focused on overall microbiome responses rather than sex-specific differences.

The gut microbiomes were collected and biobanked according to our previously validated workflow^20^. Briefly, fecal samples were collected off-site from participants and subsequently processed in the laboratory under anaerobic conditions to prepare a 20% (w/v) fecal slurry in a stabilization buffer consisting of 1× phosphate-buffered saline (PBS), 10% (v/v) glycerol, and 1 mg/mL L-cysteine. Samples were aliquoted in 15 mL Falcon tubes and stored at -80 °C until culturing. We have previously validated that the live microbiota biobanking approach can reproduce the metaproteomic responses of fresh stool samples^20^.

### Key resources

Key resources used in this study, including reagents for microbiota culturing, metaproteomics analyses, experimental consumables, instruments and software are summarized in Supplementary Table S1.

### RapidAIM 2.0 assay

#### Drug plate preparation

For quality control (QC), each plate included triplicate wells for DMSO (negative control) and triplicate wells for kestose (positive control). Blank wells contained only media and were not inoculated; they were designed to be tested for protein concentration to confirm the absence of contamination. Information of the tested compounds is given in Supplementary Data S1. For most compounds, 10 mM stocks in DMSO were made. For compounds without exact molecular weights (e.g. mixtures of polymers), 1 mg/ml stocks were prepared. For compounds dissolvable in ddH_2_O but not in DMSO, the stocks were prepared in ddH_2_O. RapidAIM 2.0 assay plates were next prepared by adding 50 μL of each stock solution in the well designated to the specific compound to achieve 500 μM (or 50 μg/mL for compounds without exact molecular weights) concentration in the final culture system. Compounds were distributed randomly across four plates. For negative control wells, 50 μL of DMSO was used. For compound stocks prepared with ddH_2_O, 50 μL of DMSO was added to control the DMSO effect. The plates were stored at -20 °C before the culturing experiment following RapidAIM 2.0 workflow.

#### Microbiome culturing and drug treatment

Culture media was optimized and validated in our previous study^41^, which includes the following composition: 2.0 g/L peptone water, 2.0 g/L yeast extract, 0.5 g/L L-cysteine hydrochloride, 2 mL/L Tween 80, 5 mg/L hemin, 10 μL/L vitamin K1, 1.0 g/L NaCl, 0.4 g/L K_2_HPO_4_, 0.4 g/L KH_2_PO_4_, 0.1 g/L MgSO_4_⋅7H_2_O, 0.1 g/L CaCl_2_⋅2H_2_O, 4.0 g/L NaHCO_3_, 4.0 g/L porcine gastric mucin, 0.25 g/L sodium cholate and 0.25 g/L sodium chenodeoxycholate. RapidAIM 2.0 culturing experiments were performed in an anaerobic chamber whose gas contained 5% H_2_, 5% CO_2_ and 90% N_2_. The drug plate containing 50 µL DMSO and a compound in each treatment well was added with 1 mL culture media and 100 µL fecal slurries as inoculum. Samples were mixed sufficiently using multi-channel pipettes before covering the wells tightly with a perforated silicone gel mat and sealed around using lab tape to prevent popping up due to gas production in the culture. The deepwell plates were shaken at 500 rpm on an orbital shaker for 48 hours at 37 °C.

#### Microbial cell washing

After culturing, culture plates were centrifuged at 3,000 g for 45 minutes at 4 °C to collect the microbial cells. The cell pellets were then washed three times with 1 mL of cold PBS. In the final wash cycle, the pellets were resuspended in an additional 1 mL of cold PBS and briefly centrifuged at 300 g for 5 minutes at 4 °C to clear debris. The resulting supernatant was carefully transferred to a new deep well plate and subjected to a final centrifugation at 3,000 g for 45 minutes at 4 °C. The purified cell pellets were subsequently stored at -80 °C until needed for cell lysis.

#### Microbial cell lysis and protein double-precipitation

Proteins were then extracted from the cells using ultra-sonication and were purified using a double-precipitation procedure. Briefly, microbial cell pellets were lysed with 150 μL of lysis buffer (8 M urea and 4% sodium dodecyl sulfate (SDS) in 100 mM Tris-HCl, pH = 8.0) per well using a 96-channel liquid handler. The lysates underwent sonication using Qsonica Q700 at 10 kHz, alternating 10 seconds on and 10 seconds off for a total of 10 minutes, with cooling at 8 °C to prevent overheating. The sonicated lysates were precitipated overnight in ice-cold protein precipitation solution (50%:50%:0.1% (v:v:v) of acetone:ethanol:acetic acid) at -20 °C. Following the first precipitation, the proteins were pelleted and resuspended in 150 μL of protein resuspension buffer (6 M urea in 100 mM Tris-HCl, pH = 8.0) and subjected to a second precipitation under the same condition.

#### Protein digestion

Protein concentrations of the DMSO control samples were measured using the DC Protein Assay kit according to the manufacturer’s instructions. An equal volume approach was applied similar to described previously^4^, ensuring comparable protein biomass measurements between samples: DMSO control samples were diluted to 1 μg/μL with the resuspension buffer (6 M urea in 100 mM Tris-HCl, pH 8), and the same volume of the resuspension buffer was added to other sample wells. Protein digestion was performed using an automated liquid handler (Hamilton Nimbus 96) following standard workflows of reduction, alkylation, and overnight tryptic digestion as described in the RapidAIM 2.0 protocol^22^.

#### Protein desalting and TMT labeling

After digestion, automated desalting of protein lysate was performed using reverse-phase (RP) desalting columns (IMCS, 04T-H6R05-1-10-96). A 240 µL aliquot of the 800 µL desalting elute was used for 96-well format TMT 11plex™-labeling following an economic and robust approach that was previously described^21^. Briefly, 11 different tandem mass tags (TMT) reagents are first resuspended in acetonitrile, aliquoted into precise 50 µg TMT per channel, freeze-dried, and stored at -80 °C. For peptide labeling, the peptide aliquots (dissolved in 100 mM TEAB with 20% acetonitrile) are incubated in the TMT reagent plates at 25 °C for 1-2 hours. Post-incubation, the reaction is quenched, and the pH adjusted to 2-3. Finally, the labeled peptides are combined, desalted, and freeze-dried for further analysis.

### LC-MS/MS analysis

#### TMT-labelled samples

LC-MS/MS analyses of TMT-labeled samples were performed using an UltiMate 3000 RSLCnano system coupled to an Orbitrap Exploris 480 mass spectrometer (ThermoFisher Scientific Inc.). Peptides were separated using a 75 μm × 50 cm analytical column packed with reverse phase beads (1.9 μm/120-Å ReproSil-Pur C18 resin, Dr. Maisch HPLC GmbH), applying a two-hour gradient from 5%∼35% of buffer B (0.1% (v/v) formic acid with 80% (v/v) acetonitrile) ratio with buffer A being 0.1% formic acid, at a flow rate of 300 nL/min. Full MS scans were acquired in the Orbitrap at a resolution of 120,000 for 350-1,200 m/z. Data were acquired in a data-dependent mode with a cycle time of 2 seconds between master scans. MS/MS scans were performed using higher-energy collisional dissociation (HCD) at a resolution of 45,000, with a collision energy of 30%. The dynamic exclusion was set to 60 seconds after a single count.

#### Label-free samples

LC-MS/MS analyses of label-free samples were also performed using the UltiMate 3000 RSLCnano and the Orbitrap Exploris 480. Peptides were separated using a tip column (75 μm inner diameter × 15 cm) packed with reverse phase beads (3 μm/120 Å ReproSil-Pur C18 resin, Dr. Maisch HPLC GmbH), applying a one-hour gradient from 5%∼ 35% of buffer B (0.1% (v/v) formic acid with 80% (v/v) acetonitrile) ratio, with buffer A being 0.1% formic acid, at a flow rate of 300 nL/min. Full MS scans were acquired in the Orbitrap at a resolution of 60,000 for 350-1,200 m/z by data-dependent MS/MS scan of the 15 most intense ions. MS/MS scans were performed using higher-energy collisional dissociation (HCD) at a resolution of 15,000, with a collision energy of 30%. The dynamic exclusion repeat count was set to one, and the repeat exclusion duration was set to 20 seconds.

### Metagenomics analysis

Fecal slurries of different individual microbiomes were first subjected to whole metagenomic DNA extraction using the FastDNA Spin Kit (MP Biomedicals, 116540600). Quantification of DNA contents was performed using the Qubit High Sensitivity dsDNA Assay Kit, following the manufacturer’s instructions^42^. Metagenomic DNA sequencing was performed with Illumina NovaSeq 6000 platform at Génome Québec CES using PCR-free library preparation. A total of 2.58B reads were sequenced for six samples with an average of 430M (±70M) per individual microbiome. The generated fastq sequences were preprocessed using fastp v.0.23.1^43^ to remove adaptors and quality filter the reads with default parameters. Human and phiX reads were then removed by mapping the reads to a non-redundant version of the Genome Reference Consortium Human Build 38 (GRCh38hg38; RefSeq: GCF_000001405.39) and phiX reference genomes using the Kraken2 v.2.1.2 package^44^. The quality-filtered paired-end reads were then error-corrected using bbcms v38.34 from the BBTools^45^ with default parameters. An average of 420M (±69M) reads were obtained after these steps. Metagenome assembly of the error-corrected and quality-filtered non-human reads was performed using MEGAHIT v1.2.9^46^ using the --presets meta-large --min-contig-len 1000 parameters. A total of 640,099 contigs with lengths >1,000 bp were individually assembled, with an average of 106,683 (±32,133) contigs per individual microbiome. Metagenomic binning was performed using SemiBin v1.5.1^47^ followed by quality filtering (completeness >50%) with DAS Tool v1.1.4^48^, the high-quality bins obtained through this process were considered as metagenome-assembled genomes (MAGs)^23^. Species-and genus-level richness and shared taxa across metagenomes are shown as Supplementary Figure S8.

The assembled contigs were then annotated using the PROkaryotic Dynamic programming Gene-finding ALgorithm (Prodigal) v2.6.3^49^ to predict open reading frames (ORF). The contigs were translated into amino acid sequences using the anonymous gene prediction mode (prodigal -p meta) and default parameters and annotated a total of 3.8M protein-coding genes. Lastly, the outputs were compiled into a single FASTA file and used as the metagenome-inferred protein database for the metaproteomic search.

### Database search strategy

The initial compound screening was performed using TMT-labeled data analyzed with MetaLab (Version 2.3.0)^50^, which implements an automated spectral clustering algorithm to group similar MS/MS spectra prior to database searching against the IGC reference database^51^. The samples selected from TMT-based results were re-analyzed by label-free LC-MS/MS quantification. Since reference catalog-based databases often lack microbiome-specific peptides^52^, we developed a bioinformatic workflow for sample set specific database generation (Supplementary Figure S1a). Briefly, metagenomic prodigal sequences (in FASTA format) from each individual microbiome were used to construct reduced databases of each subset of individual microbiome’s LFQ LC-MS/MS raw files following the MetaPro-IQ workflow^26^. First, databases were reduced from two-step X!Tandem search to remove non-expressed sequences based on actual MS data of each subject’s raw file subsets. Next, reduced subject-specific databases were merged and subjected to redundancy removal at 99% using CD-HIT. The resulting FASTA file containing 491,506 sequences was used for database search of LC-MS/MS label-free files using MaxQuant (version 1.5.2.8), as well as revisiting the TMT dataset’s database search using MaxQuant (version 2.0.3.0). Both LFQ and TMT searches applied carbamidomethylation as a fixed modification (C) and variable modifications including acetylation at the protein N-terminus and oxidation of methionine, with a fragment ion tolerance of 20 ppm. The false discovery rates (FDR) for peptide spectrum matches (PSM) and proteins were set to 0.01, with a minimum peptide length of 7. Match between runs were enabled with matching time window of 0.7 minutes and alignment time window of 20 minutes. To specifically investigate compound effects on protein expressions related to antimicrobial resistance, we conducted a focused search against the full CARD database^35^ (downloaded December 27^th^, 2022) using MaxQuant 1.5.2.8, with the same search parameters as above.

### Data preprocessing

For the TMT dataset, proteinGroup.txt was normalized using MSstatsTMT^53^ with the “msstats” option selected, and perform estimate size factors on matrix using Deseq2 R package, the processing resulted in 19,318 proteinGroups across all TMT samples. A Q15 criteria was applied to remove highly random sparse features, and SpectroFM was applied for data imputation, resulting in a matrix of 6085 proteinGroups.

For the LFQ dataset, directLFQ^27^ (GUI MacOS version 0.2.8) was used for efficient label-free quantification, followed by protein inference using a custom in-house script (available at https://github.com/ningzhibin/protein-Inference). Razor and unique peptides were considered for protein quantification. Dataset was further harmonized by individual microbiome batch using the R package HarmonizR^28^, which implements the ComBat algorithm while appropriately handling missing values^28^.

For the ARP database search result, protein intensities in the proteinGroups file were filtered to remove potential contaminants and reverse sequences, then all protein intensities in each sample were summed as ARP intensities. The ARP ratio in each sample was then calculated by dividing the ARP intensities by the total protein intensities in the LFQ search result.

### Taxonomic and functional annotation

Taxonomic annotation was performed in MetaLab 2.3.0^50^ by searching the peptide sequences against the pep2taxa database in MetaLab. Taxonomic annotations with three or more unique peptides were considered confident, following the conservative criteria used in previous studies^50,54^. eggNOG mapper (version 2.1.12)^55^ was used for functional annotation of protein sequences, and COG functional enrichment of selected eggNOG matches was performed using the Enrichment Analysis tool of iMetaShiny^56^.

### Downstream data analysis

#### Compound structure clustering

Clustering of compounds was performed using the fingerprint, and rcdk R packages following a method developed by Voicu et al.^57^ Briefly, The SMILES (Simplified Molecular Input Line Entry System) notation for each compound (excluding those without a defined chemical structure) was parsed using the parse.smiles() function from the rcdk package, then compound fingerprints were obtained by the get.fingerprint() function of the fingerprint package using type “extended” as suggested by Voicu et al.^57^ Next, fp.sim.matrix() function was used to calculates the similarity matrix for the set of compound fingerprints by the Tanimoto coefficient. Finally, the compound structure similarity matrix was passed to the pvclust function, which performed hierarchical clustering and generated 100 bootstrap samples to evaluate the stability of the clustering results.

#### Kernel density estimation (KDE)

Kernel density estimation (KDE) of the two-dimensional ARP versus *nFR_p_* landscape was performed using the kde2d() function of R package MASS with n resolution set as 100, and 3D surface plot of the KDE results was visualized by R package plotly.

#### Feature set comparison

Comparison of shared features between different groups were performed using upset plot wrapped by the “Sets Explorer” in iMetaShiny^56^ (https://shiny2.imetalab.ca/shiny/rstudio/SetExplorer/).

#### Principal component analysis

For principal component analysis of metaproteomic drug responses, protein intensity data was first log10 transformed, then normalized using the estimateSizeFactorsForMatrix() function of the R package DeSeq2. Next, the pca() function of the pcaMethods package was used to perform PCA analysis using the nonlinear estimation by the NIPALS algorithm, allowing for the presence of missing values^29^.

#### Finding microbial homologs of human protein drug targets

Human drug targets of our selected drugs were extracted from the comprehensive map of molecular drug targets (NIHMS80822-supplement-Supplementary_Information_S2.xlsx table in the referred paper) summarized by Santos et al.^30^ To identify homologs of human protein drug targets in the tested microbiomes, we employed a profile Hidden Markov Model (HMM)-based approach using the HMMER suite (v3.3). For each target protein, *hmm* and *seed* files were downloaded from the Pfam database^58^ (https://www.ebi.ac.uk/interpro/), then hmmsearch^31^ was used to perform microbial homolog sequence search by protein alignment/profile-HMM against our metaproteomic sequence database of this dataset.

#### Functional redundancy calculation

Within-sample taxonomic diversity (TD) and functional redundancy (*FR_p_*) of each metaproteome was calculated using the label-free quantified dataset following our previously developed approach^25^. Briefly, the proteome-content network of each individual metaproteome data was generated according to pep2taxa-based taxonomic annotation and eggNOG-based Clusters of Orthologous Genes (COG) annotations. Genus-level biomass contributions were estimated by genus-specific peptide intensities in pep2taxa-based taxonomic annotation. Then, taxonomic diversity (TD) and functional redundancy normalized by TD (*nFP_p_*) was calculated using the python codes provided by our previous research^25^ (https://github.com/yvonnelee1988/Metaproteome_FRp).

#### Differential protein analysis

Differential protein expression between DMSO and the seven neurological drugs (amoxapine, duloxetine, fluoxetine, fluphenazine, fluvoxamine, paroxetine and sertraline) was analyzed using the Differential Protein Analyzer of iMetaLab Suite^56^. The functional annotation and taxonomic representatives of the resulting proteins were annotated by eggNOG mapper and visualized using Sanky plot with geom_alluvium() function of ggplot2.

#### PhyloFunc analysis and community state types (CST) clustering

The phylogenetic tree was constructed from the deep metagenomics dataset using IQ-TREE2 (http://www.iqtree.org/), the tree inference included 1000 replicates for the SH-like approximate likelihood ratio test (aLRT) and 1000 ultrafast bootstrap replicates. PhyloFunc was then computed with the phylogenetic tree and the original directLFQ-quantified protein group table, with taxonomy information annotated by referencing the metagenome-assembled genomes (MAGs), and functional annotation performed by eggNOG mapper^55^. The Python package *phylofunc* ^23^ was then used to compute functional distances matrices across metaproteomics samples.

#### Other analyses and visualizations

Correlation analysis was performed using cor.test() function in the stats package. Scatter plots with aligned density plots were created by ggscatterhist() function in the ggpubr package. Partitioning around medoids analyses were performed using pam() function in the cluster package. Taxonomic heatmap and clustering was performed using the pheatmap package.

### Computation of the generalized Lotka-Volterra (gLV) model

The gLV competition model, considering between-cluster interactions can be written as follows:

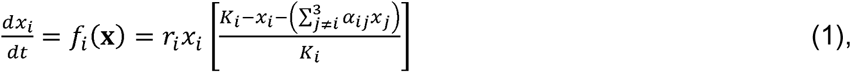

where, *i* and *j* represent *i*^th^ and *j*^th^ microbial cluster, respectively. In our case, 1 ≤ *i*≠*j* ≤ 3. **x**=[*x_1_*, *x_2_*, *x_3_*] represents the population size of each microbial cluster. The parameters *r_i_*, *K_i_* and *α_ij_* represent the intrinsic growth rate of the *i*^th^ microbial cluster, environmental carrying capacity of the *i*^th^ microbial cluster, and competition coefficient from the *j*^th^ to the *i*^th^ microbial cluster, respectively. We note that the stable state is dependent on model parameters *r_i_*, *K_i_* and *α_ij_* rather than the initial population size of each microbial cluster. Based on the gLV competition model considering between-cluster interactions given by equation (1), we can calculate population sizes at the equilibrium state where 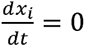. Therefore, at equilibrium, we have:

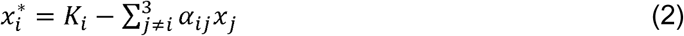

Based on Lyapunov’s indirect method, the Jacobian matrix **J** of the gLV model at the equilibrium point **x***\**=[*x*_1_*, *x*_2_*, *x*_3_*] can be obtained. By linearizing equation (1) around **x***\** and its neighborhood, it is then transformed into a linear dynamic model:

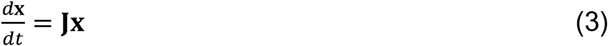

Where,

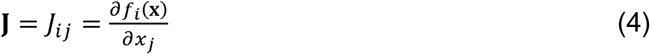

If all eigenvalues of the matrix **J** lie in the left half of the complex plane, then we achieve a stable equilibrium point of the gLV model.

Ordinary equation of the gLV model was solved using scipy.integrate function odeint() in Jupyter Notebook running Python 3. And visualization of model dynamics was performed using matplotlib.pyplot.plot().

### Functional redundancy enhancement to combat ARP response

An additional RapidAIM test was performed to investigate the impact of enhanced functional redundancy on mitigating drug-induced ARP within microbial communities. First, to evaluate the impact of FOS on functional redundancy, three additional individual microbiomes were initially screened for their response of *nFR_p_* to 10 mg/mL of FOS treatment. The microbiome exhibiting a high increase in *nFR_p_* was selected for further analysis. It was subjected to 250 µM sertraline, 10 mg/mL FOS, or 250 µM sertraline + 10 mg/mL FOS in 25 µL DMSO, and blank (25 µL DMSO only) treatments for 48 hours of culturing, followed by metaproteomic analyses to assess changes in functional redundancy and ARP levels in different groups. Technical triplicates were performed for each group.

### Statistics and Reproducibility

Estimation of sample size was based on practical considerations of large-scale drug screening (312 compounds + 6 control samples per biological replicate), which necessitated minimizing the number of biological replicates per condition. Using an effect size of Cohen’s d = 1.2 (as confirmed by post hoc analysis of the DMSO vs Sertraline comparison), with a significance level of 0.05 and a statistical power threshold of 0.7, the minimum required sample size was estimated to be six biological replicates per group. All data were retained for analysis unless missingness was caused by technical issues during sample processing. One DMSO control sample from subject V47 was lost during the mass spectrometry acquisition step. In addition, sample V52_P1E3 yielded only 290 quantified values after database searching and was excluded due to insufficient data quality. The distribution of compounds across 96-well plates was randomized.

Due to the nature of automated proteomics sample processing, the downstream handling steps were not randomized; however, for each individual, LC–MS/MS sample injection order was randomized. Although the investigators were not blinded to the allocation during experiments and outcome assessment, the automated sample processing ensured that experimental and control groups were handled in a uniform and unbiased manner.

## DATA AVAILABILITY

All metaproteomics data sets reported in this study have been deposited in the ProteomeXchange database through the PRIDE repository under accession codes PXD056930 [https://www.ebi.ac.uk/pride/archive/projects/PXD056930], PXD056888[https://www.ebi.ac.uk/pride/archive/projects/PXD056888], PXD056968[https://www.ebi.ac.uk/pride/archive/projects/PXD056968]. The metagenomics data used in this study are deposited in the NCBI Sequence Read Archive (SRA) with the accession numbers SRR30894461[https://www.ncbi.nlm.nih.gov/sra/?term=SRR30894461], SRR30894462[https://www.ncbi.nlm.nih.gov/sra/?term=SRR30894462], SRR30894463[https://www.ncbi.nlm.nih.gov/sra/?term=SRR30894463], SRR30894464[https://www.ncbi.nlm.nih.gov/sra/?term=SRR30894464], SRR30894465[https://www.ncbi.nlm.nih.gov/sra/?term=SRR30894465] and SRR29021656[https://www.ncbi.nlm.nih.gov/sra/?term=SRR29021656].

## CODE AVAILABILITY

Code and related data to reproduce the analyses in this study has been published in figshare through DOI: 10.6084/m9.figshare.29816780. In addition, the source code of the Shiny app is made publicly available via GitHub at https://github.com/yvonnelee1988/MPR_Viz and Zendo at DOI: 10.5281/zenodo.16990829. Any additional information required to reanalyze the data reported in this paper is available from the lead contact (Daniel Figeys) upon request.

## ACKNOWLEDGMENTS

L.L acknowledges funding from National Natural Science Foundation of China (grant 32370050), and the State Key Laboratory of Medical Proteomics (SKLP-Y202402). D.F. acknowledges funding from Genome Canada, the Province of Ontario and NSERC. The authors would like to thank Dr. Dawei Hu for his advice and help on the gLV model, and Dr. Yangyu Liu for his advice on functional redundancy analysis. The authors also acknowledge technical support platforms and staffs at the Faculty of Medicine, University of Ottawa and National Center for Protein Sciences (Beijing) for providing essential resources and assistance.

## AUTHOR CONTRIBUTIONS

D.F. and L.L. designed and supervised the study, and wrote the manuscript; M.H., C.M.A.S., L.L., Xu Z. and D.F. selected compounds for this study. J.M., Xu Z., K.W. and H.Q. recruited healthy volunteers, collected individual fecal samples, collected written consent and biobanked the samples. M.H. and L.L. prepared the therapeutic compound library. L.L. established, validated, and performed the RapidAIM in vitro culturing and metaproteomic sample processing workflows, during which process, J.M., K.W., M.H. and H.Q. assisted with microbiome culturing and microbial cell washing. L.L. and Xin Z. performed the functional redundancy analyses. Z.N. optimized LC-MS/MS parameters, both Z.N. and L.L. performed the metaproteomic LC-MS/MS analysis and tracked LC-MS/MS performance. L.L., K.C. and C.M.A.S. performed metaproteomic database search. J.B. and A.S. performed metagenomic sample processing and sent samples for Illumina NovaSeq analysis. J.M.S. performed metagenomic data analysis and created metagenomics-based protein databases. L.L. and L.W. performed the PhyloFunc analysis. L.L. and C.M.A.S curated the data and performed TMT metaproteomics data analysis and compound structure clustering. L.L. performed all label-free metaproteomics down-stream data analysis. All authors contributed to the writing, editing, and proofreading of the manuscript.

## COMPETING INTERESTS STATEMENT

D.F. and A.S. are co-founders of MedBiome Inc., a microbiome nutrition and therapeutic company. C.M.A.S. is currently a staff at Recursion Pharmaceuticals, and K.W. is currently a staff at Carleton University, the presented contribution in this manuscript was made during their employment at the University of Ottawa. Other co-authors declare no conflict of interest.

## Notes

https://shiny.imetalab.ca/MPR_Viz/

### Summary of Updates

This version has been revised according to the reviewers' and editors' comments during the peer-review process at Nature Communications.

