## Supplementary Table S1 for "Systematic Metaproteomics Mapping Reveals Functional and Ecological Landscapes of Ex Vivo Human Gut Microbiota Responses to Therapeutic Drugs"

### Supplementary Table S1. Key resource table

| REAGENT or RESOURCE | SOURCE | DENTIFIER |
|---|---|---|
| RapidAIM 2.0 Assay – Microbiota culturing | | |
| Potassium phosphate monobasic | Millipore-Sigma - Sigma-Aldrich | P5655 |
| Potassium phosphate dibasic | Millipore-Sigma - Supelco | PX1570-1 |
| Sodium chloride | Calbiochem - OmniPur® | 7710 |
| Magnesium sulfate heptahydrate | Millipore-Sigma - Sigma-Aldrich | 230391 |
| Calcium chloride, anhydrous | Millipore-Sigma - Sigma-Aldrich | C5670 |
| Tween® 80 | Millipore-Sigma - Sigma-Aldrich | P4780 |
| Sodium cholate hydrate | Millipore-Sigma - Sigma-Aldrich | C9282 |
| Sodium chenodeoxycholate | Millipore-Sigma - Sigma-Aldrich | C8261 |
| Peptone water | Millipore-Sigma - Sigma-Aldrich | 70179 |
| Bacto™ yeast extract | Becton, Dickinson and Company | 212750 |
| Sodium bicarbonate | Millipore-Sigma - Supelco | SX0320 |
| L-cysteine | Millipore-Sigma - Sigma-Aldrich | C7352 |
| 1-Kestose | TCI America | K0032 |
| Hydrochloric acid | Fisher Chemical | A144S |
| RapidAIM 2.0 Assay – Metaproteomic sample processing | | |
| Phosphate buffered saline, 10X solution | Fisher BioReagents | BP399-4 |
| Urea | Millipore-Sigma - Sigma-Aldrich | U5378 |
| Tris (hydroxymethyl)aminomethane | Calbiochem - OmniPur® | 9230 |
| Sodium dodecyl sulfate | Millipore-Sigma - Sigma-Aldrich | L3771 |
| Acetone | Millipore-Sigma - Sigma-Aldrich | 179124 |
| Acetic acid, glacial | Fisher Chemical | A38-212 |
| Acetonitrile | Millipore-Sigma - Sigma-Aldrich | 34851 |
| Ethyl alcohol, anhydrous | Commercial Alcohols | P016EAAN |
| Formic acid | Millipore-Sigma - Sigma-Aldrich | F0507 |
| cOmplete™ protease inhibitor cocktail | Millipore-Sigma - Roche | 04693116001 |
| Dithiothreitol | Millipore-Sigma - Sigma-Aldrich | 43815 |
| Iodoacetamide | Millipore-Sigma - Sigma-Aldrich | I1149 |
| Trypsin | Worthington Biochemical | L5003740 |
| DC Protein Assay Reagents A, B and S | Bio-Rad Laboratories | 5000113, 5000114 and 5000115 |
| RapidAIM 2.0 Assay – TMT labelling | | |
| TMT10plex Isobaric Label Reagent Set plus TMT11-131C Label Reagent | Thermo Scientific | A34808 |
| 50% hydroxylamine (HOA) for TMT experiments | Thermo Scientific | 90115 |
| 1M triethylammonium bicarbonate (TEAB) for TMT experiments | Thermo Scientific | 90114 |
| Pierce™ quantitative colorimetric peptide assay | Sigma-Aldrich | 23275 |
| RapidAIM 2.0 Assay – Consumables | | |
| Culture plate | Sigma-Aldrich | CLS3960 |
| Culture plate lid | Thermo Scientific | AXYAM2MLSQ |
| Lysis plate | Thermo Scientific | AB-0600 |
| Lysis plate lid | Sigma-Aldrich | AB-0783 |
| Precipitation plate | Sigma-Aldrich | CLS4413 |
| Precipitation plate lid | Thermo Scientific | CLS4418 |
| Elution plate | Thermo Scientific | AB-0859 |
| Elution plate lid | Thermo Scientific | 278616 |
| TMT plate | Thermo Scientific | 249944 |
| TMT plate lid | Axygen™ | 276002 |
| Reservoir | Eppendorf | RESSW96HP |
| RapidAIM 2.0 Assay – Automation | | |

| | | |
|---|---|---|
| Automated 96-channel liquid handling platform | Hamilton | OPP041219 |
| Desalting tip columns | IMCS | 04T-H6R05-1-10-96 |
| RapidAIM 2.0 Assay – LC-MS/MS Analysis | | |
| UltiMate 3000 RSLCnano system | Thermo Fisher Scientific | ULTIM3000RSLCNANO |
| Orbitrap Exploris 480 mass spectrometer | Thermo Fisher Scientific | BRE725533 |
| Metagenomic Analysis | | |
| FastDNA Spin Kit | MP Biomedicals | 116540600 |
| Qubit High Sensitivity dsDNA Assay Kit | Thermo Fisher Scientific | Q32851 |
| Illumina NovaSeq 6000 | Illumina | NovaSeq 6000 |
| Software and bioinformatic tools | | |
| MetaLab | https://imetalab.ca | Version 2.3.0 |
| MaxQuant | https://www.maxquant.org/ | 1.5.2.8 |
| X!Tandem | https://www.thegpm.org/TANDEM/ | Version 2015.12.15.2 |
| directLFQ[27] | https://github.com/MannLabs/directlfq | MacOS version 0.2.8 |
| eggNOG mapper | http://eggnog-mapper.embl.de/ | Version 2.1.12 |
| fastp | https://github.com/OpenGene/fastp | Version 0.23.1 |
| Kraken2 | https://github.com/DerrickWood/kraken2 | Version 2.1.2 |
| Genome Reference Consortium Human Build 38 | Genome Reference Consortium | GRCh38hg38 |
| phiX reference genomes | NCBI | phiX174 |
| bbcms | BBMap suite | Version 38.34 |
| MEGAHIT | https://github.com/voutcn/megahit | Version 1.2.9 |
| Prodigal | https://github.com/hyattpd/Prodigal | Version 2.6.3 |
| R | R Foundation | Version 4.0.4 |
