## Supplementary Figures for "Systematic Metaproteomics Mapping Reveals Functional and Ecological Landscapes of Ex Vivo Human Gut Microbiota Responses to Therapeutic Drugs"

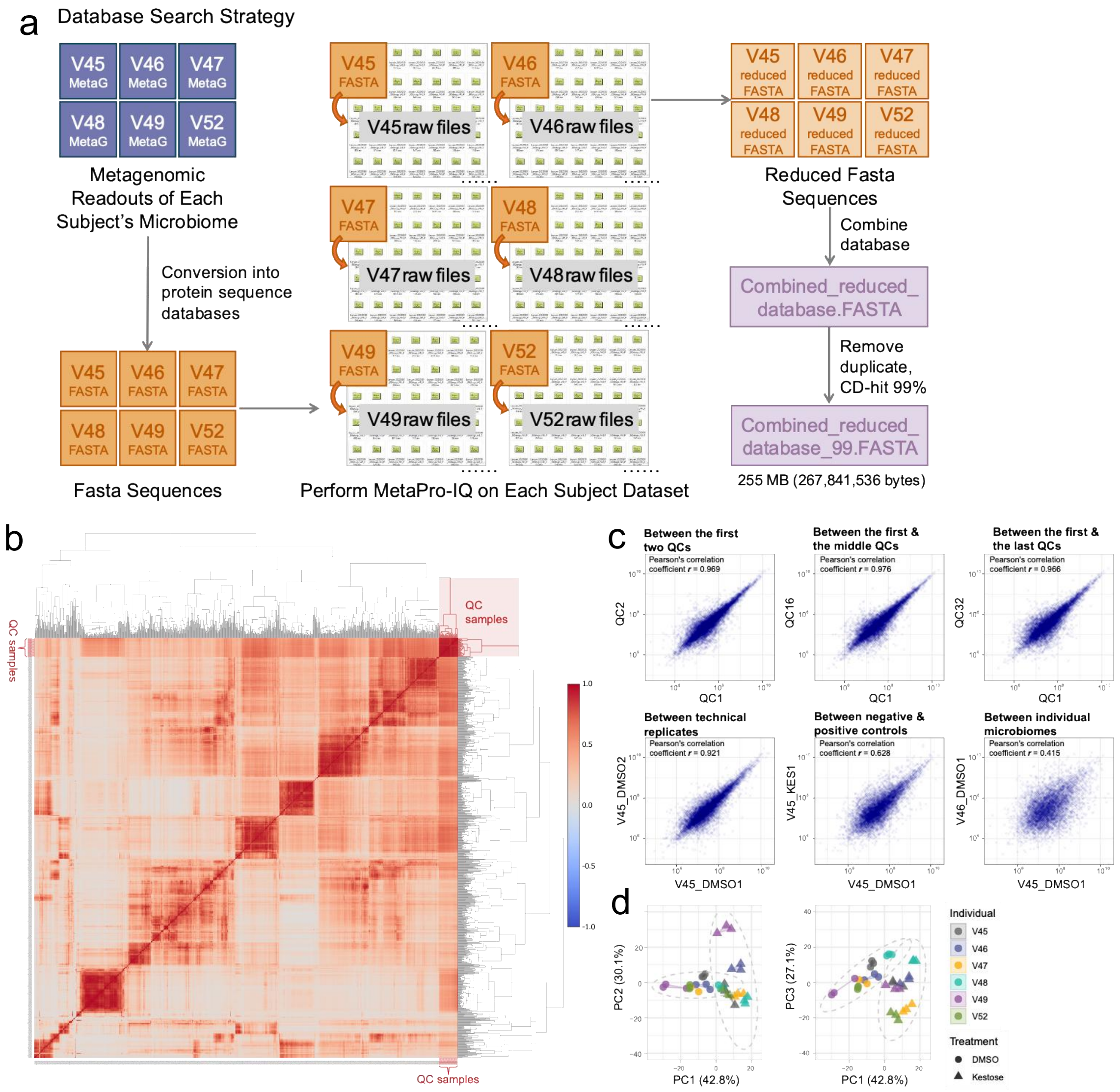


**Figure S1. Database search strategy for the Label-free LC-MS/MS dataset and the dataset quality**

**(a)** Deep metagenomic data of each subject's original microbiome was converted into protein FASTA sequences using Prodigal. All LFQ raw files corresponding to each subject's cultured microbiome were subjected to the MetaPro-IQ workflow based on each corresponding database to generate individual subset-specific reduced databases. The reduced databases were combined and duplicated proteins were removed using 99% sequence identity threshold with CD-HIT. **(b)** Label-free LC-MS/MS sample correlations. Heatmap and hierarchical clustering showing between-sample Pearson's correlation coefficients throughout the label-free LC-MS/MS sample analysis. Quality control (QC) samples were created by mixing an equal amount of negative control (DMSO) samples of the six individuals' microbiomes, one QC sample was inserted each day during the analysis, resulting in 32 QC samples throughout the run. All QC samples are highlighted in red on this figure. **(c)** Scatter plots showing reproducibility of the study as well as divergence of different groups of samples. Scatter plot and Pearson's correlation of LFQ protein intensities between the first two QC samples; between the first QC sample and the middle QC sample; between the first QC sample and the last QC sample; between one pair of technical replicates (two cultured negative control samples of V45 is shown); between the negative and positive controls (a negative control sample vs a kestose-treated microbiome, i.e. the positive control sample of V45); between the DMSO samples of V45 and V46. **(d)** PCA showing that the positive control was effective.

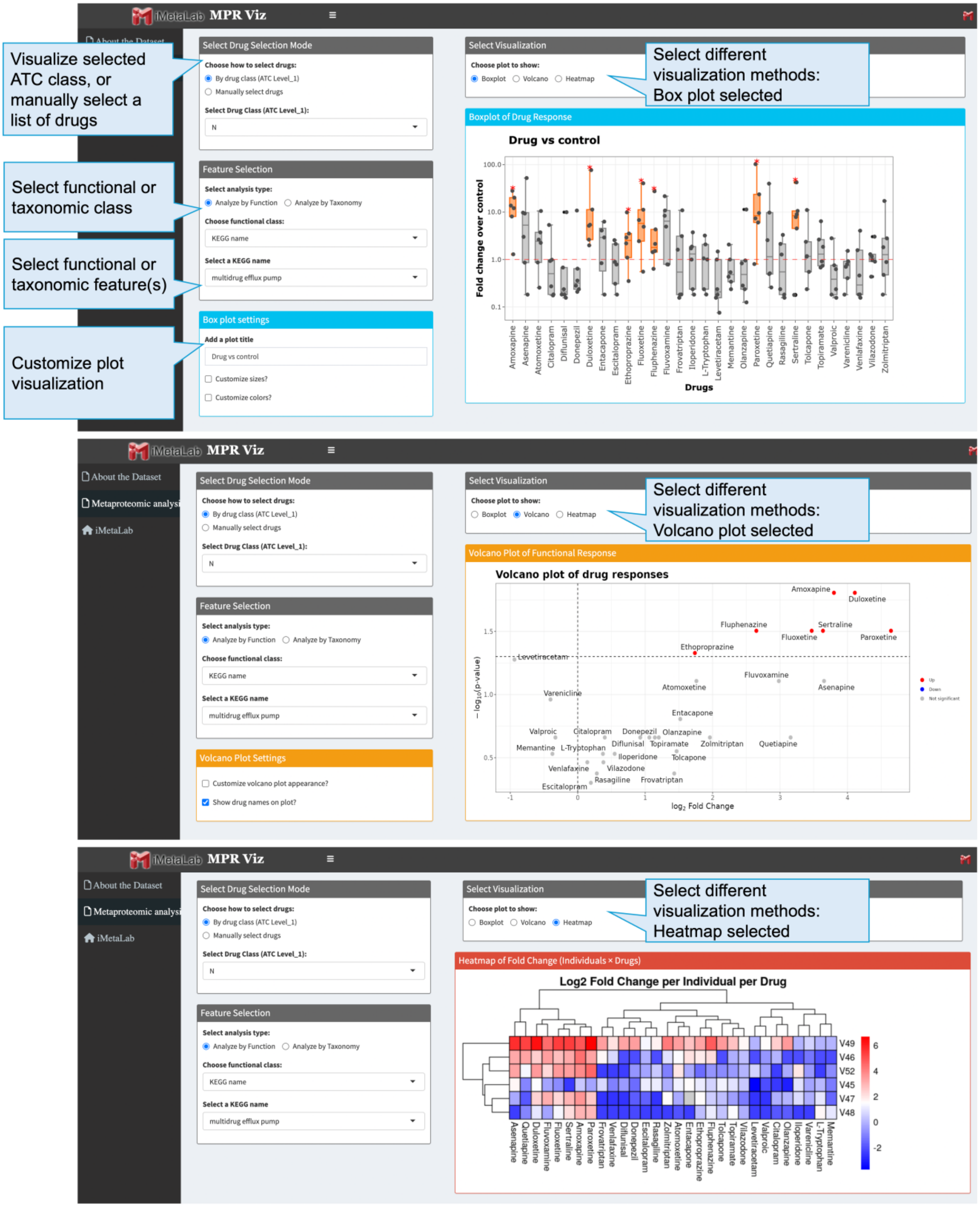
iMetaLab MPR Viz
About the Dataset
Metaproteomic analysi
iMetaLab
Visualize selected ATC class, or manually select a list of drugs
Select functional or taxonomic class
Select functional or taxonomic feature(s)
Customize plot visualization
Select different visualization methods: Box plot selected
Select different visualization methods: Volcano plot selected
Select different visualization methods: Heatmap selected
Select Drug Selection Mode
Choose how to select drugs:
By drug class (ATC Level_1)
Manually select drugs
Select Drug Class (ATC Level_1):
N
Feature Selection
Select analysis type:
Analyze by Function
Analyze by Taxonomy
Choose functional class:
KEGG name
Select a KEGG name
multidrug efflux pump
Box plot settings
Add a plot title
Drug vs control
Customize sizes?
Customize colors?
Select Visualization
Choose plot to show:
Boxplot
Volcano
Heatmap
Boxplot of Drug Response
Drug vs control
Fold change over control
Drugs
Volcano Plot Settings
Customize volcano plot appearance?
Show drug names on plot?
Volcano Plot of Functional Response
Volcano plot of drug responses
log2 Fold Change
Heatmap of Fold Change (Individuals × Drugs)
Log2 Fold Change per Individual per Drug
V49
V46
V52
V45
V47
V48

**Figure S2. Interface of the Shiny app showing an example metaproteomic response visualization.** The Shiny app is available at (https://shiny.imetalab.ca/MPR_Viz/). The analysis interface includes selection of drug category (at ATC level-1) or a customized list of compounds, functional or taxonomic annotation classes, and detailed functional or taxonomic annotation features. The plotting area is showing an example analysis on the response of microbial proteins annotated with multidrug efflux pump function to a series of N class compounds. The result data can be visualized using three different modes: box plot, volcano plot, or heatmap, providing flexible views of the responses.

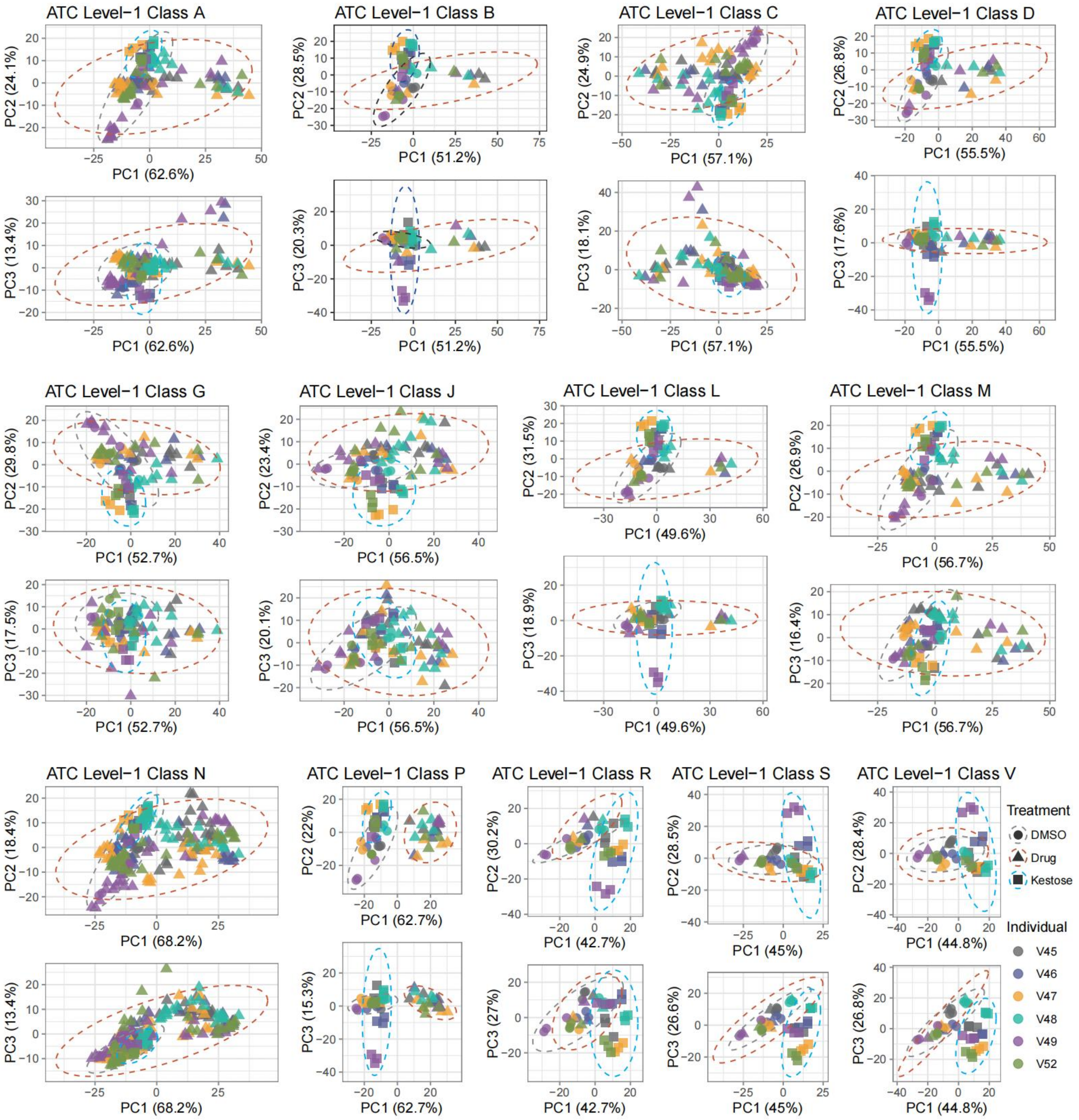


**Figure S3. PCA analysis of the metaproteomics responses by ATC level-1 classes.**

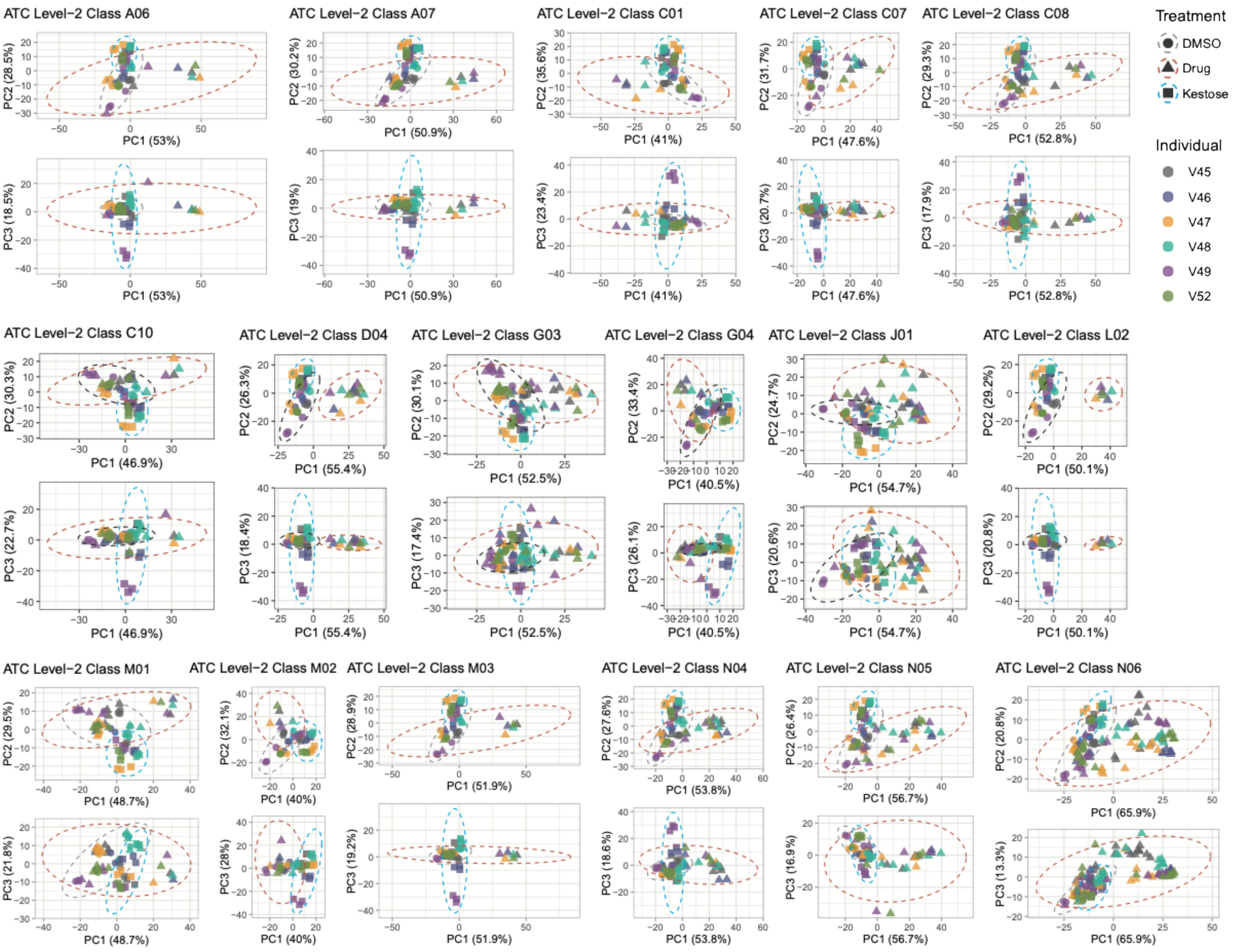


**Figure S4. Metaproteomic responses to ATC level-2 subclasses that contain MPR+ compounds.** Detailed drug names are listed in Supplementary Table S2.

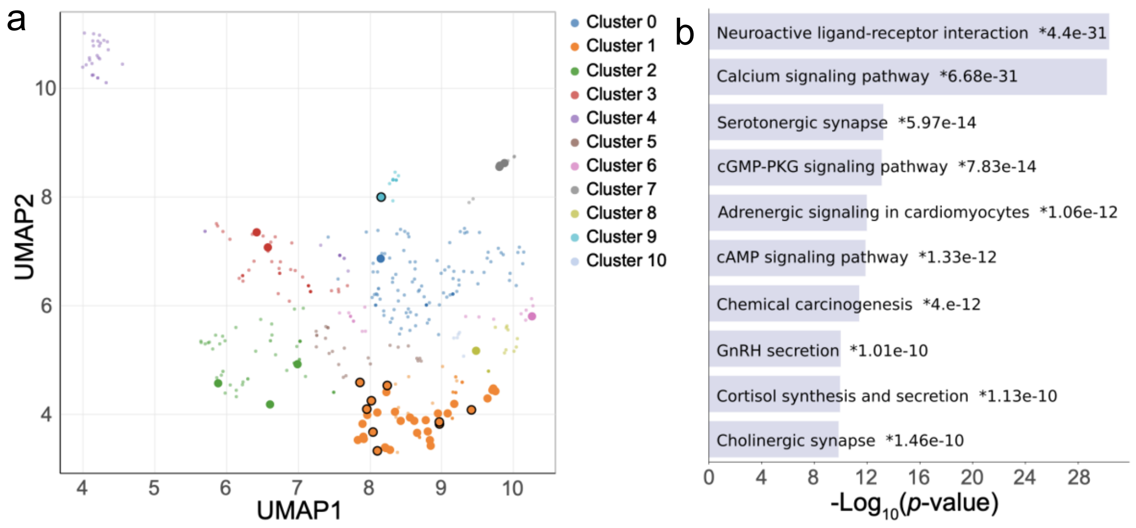


**Figure S5. Enrichment analyses of MPR+ compounds' human targets. (a)** 50 human protein targets corresponding to all MPR+ compounds were subjected to enrichment analysis against the KEGG_2021_Human gene set library. Darker and larger points, indicate more significantly enriched terms compared to genes in the library. Here we show that significantly enriched KEGG terms are concentrated in Cluster 1. **(b)** Bar plot of 10 most highly enriched terms among the human protein targets against the KEGG_2021_Human gene set database. Enrichment analyses were performed using Enrichr on the Appyter platform[59], in which the p-values are computed from a one-sided Fisher exact test to evaluate the over-representation of input gene sets against the background database, shown are raw *p*-values uncorrected for multiple hypothesis testing.

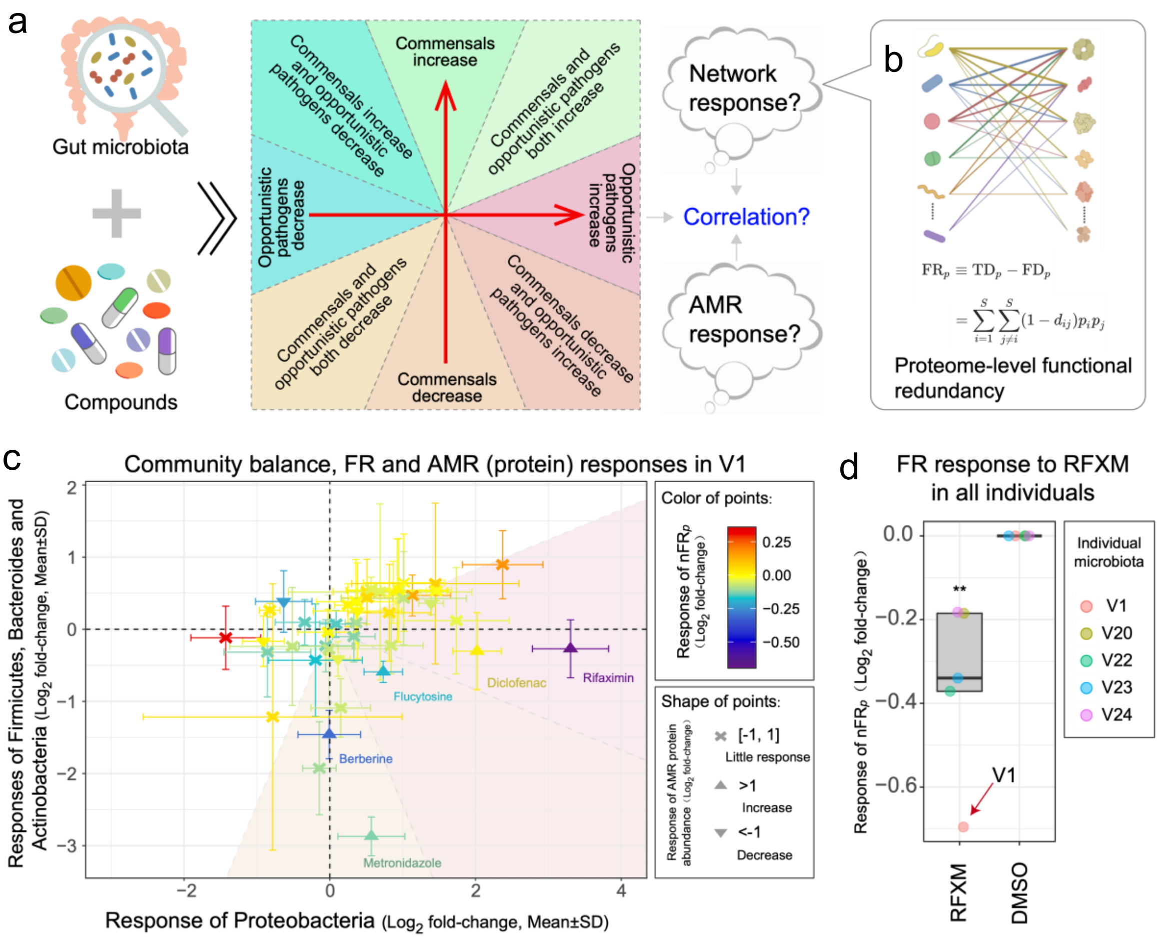


**Figure S6. Re-analysis of a previously published dataset suggesting possible connection between ARP and FR$_p$. (a)** Drug interventions on the gut microbiota can lead to changes in the balance between symbiotic bacteria and opportunistic pathogens. Therefore, we can map these changes onto a Cartesian coordinate system, dividing them into different regions (as indicated by color blocks and text annotations), to visually observe the drug response on microbiota balance. It is of interest whether samples falling into different regions correlate with the ecological properties of the community and the response of antibiotic resistance proteins. **(b)** The protein-level functional redundancy of individual microbiota samples can be quantified as described in our previous work[25]. **(c)** The normalized functional redundancy ($nFR_p$) and the response of antibiotic resistance proteins to multiple drugs in the gut microbiota of individual V1 from our previous dataset[4] was calculated and presented on the community balance map. We visualized the protein biomass along the x-axis representing the response of the Proteobacteria phylum (now referred to as Pseudomonadota in the updated NCBI taxonomy), which is rich in opportunistic pathogens, and the y-axis representing the response of the Firmicutes (Bacillota), Bacteroidetes (Bacteroidota), and Actinobacteria (Actinomycetota) phyla, which are primarily symbiotic bacteria. A $Log_2$ fold-change >1 indicates that the abundance of antibiotic resistance proteins after drug treatment is more than twice that of the DMSO control group. **(d)** In all five individuals in the previous dataset[4], the functional redundancy of their gut microbiota showed significant reductions under rifaximin (RFXM) treatment, with individual V1 exhibiting the largest reduction (indicated by the red arrow) (n=5 individual gut microbiomes). Statistical significance was observed (p = 0.00965, one-sided *t*-test), as indicated by **. The center line of box represents the median; the box limits indicate the upper and lower quartiles (Q3 and Q1); the whiskers extend to the most extreme values within 1.5× the interquartile range (IQR) from the quartiles.

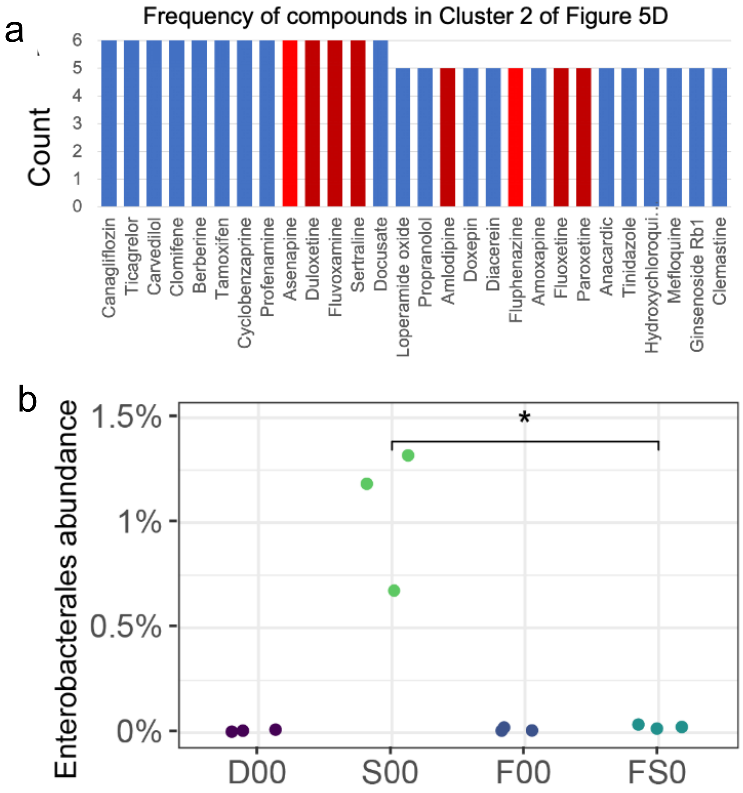


**Figure S7. Sertraline increased ARP, reduced FRp, and elevated Enterobacteriales abundance. (a)** Frequency of drugs in cluster 2 of the six ARP vs $FR_p$ plots (as in Figure 6D) showing Sertraline increased ARP and reduced $FR_p$ in all six individual microbiomes. **(b)** Sertraline increased Enterobacterales abundance, and this increase was reversed by the addition of FOS (n=3 technical replicates). Statistical significance of 0.034 (as indicated by *) was observed between Sertraline and FOS+Sertraline groups with two-sided *t*-test.

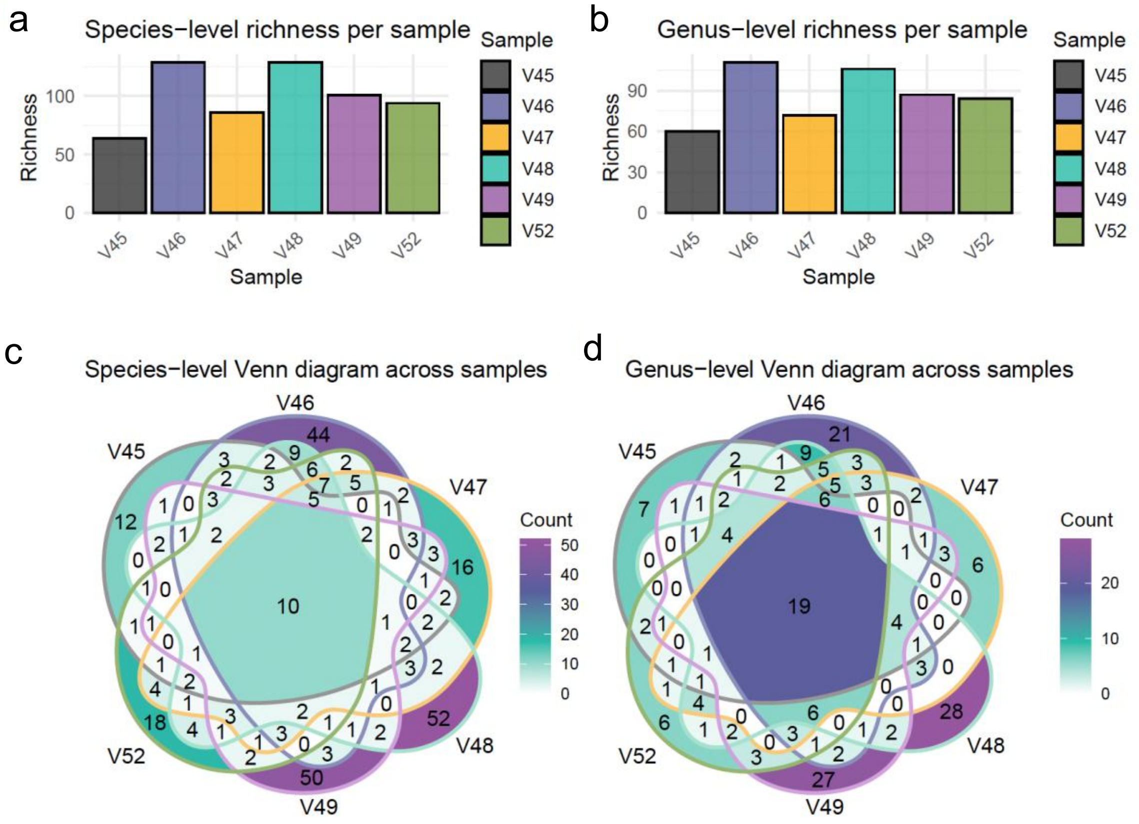


**Figure S8. Species- and genus-level richness and shared taxa across samples.** **(a–b)** Richness per sample at species and genus levels. Each bar represents one single value corresponding to each sample. **(c–d)** Venn diagrams showing shared and unique taxa across samples at species and genus levels.
